# In Vivo Biorthogonal Antibody Click for Dual Targeting and Augmented Efficacy in Cancer Treatment

**DOI:** 10.1101/2023.09.05.556426

**Authors:** Sandeep Surendra Panikar, Na-Keysha Berry, Shayla Shmuel, Nai Keltee, Patrícia M.R. Pereira

## Abstract

Antibody-drug conjugates (ADCs) have emerged as promising therapeutics for cancer treatment; however, their effectiveness has been limited by single antigen targeting, potentially leading to resistance mechanisms triggered by tumor compensatory pathways or reduced expression of the target protein. Here, we present antibody-ADC click, an approach that harnesses bioorthogonal click chemistry for *in vivo* dual receptor targeting, irrespective of the levels of the tumor’s expression of the ADC-targeting antigen. Antibody-ADC click enables targeting heterogeneity and enhances antibody internalization and drug delivery inside cancer cells, resulting in potent toxicity. We conjugated antibodies and ADCs to the bioorthogonal click moieties tetrazine (Tz) and trans-cyclooctene (TCO). Through sequential antibody administration in living biological systems, we achieved dual receptor targeting by *in vivo* clicking of antibody-TCO with antibody-Tz. We show that the clicked antibody therapy outperformed conventional ADC monotherapy or antibody combinations in preclinical models mimicking ADC-eligible, ADC-resistant, and ADC-ineligible tumors. Antibody-ADC click enables *in vivo* dual-antigen targeting without extensive antibody bioengineering, sustains tumor treatment, and enhances antibody-mediated cytotoxicity.

## INTRODUCTION

Antibody-drug conjugates (ADCs) have transformed cancer care, with over 14 FDA-approved ADCs being effective in the treatment of patients with cancer (*1*). The global ADC market is projected to reach over $15 billion USD by 2030 (*2*) attributed to the exceptional specificity of ADCs that allows targeted delivery of potent cytotoxic drugs to cells expressing the target antigen (*3–5*). However, ADCs rely on single antigen targeting, which can result in resistance if tumors activate compensatory pathways in response (*1, 6, 7*) or if the expression of the target protein is decreased (*8*)(*9*).

To overcome the limitations of single-antigen targeting by ADCs, both antibody (mAbs) combination therapies (*10–12*)(*13*) and bispecific antibodies (*14*) have been explored and approved for cancer treatments. While mAbs plus ADC combinations, targeting distinct epitopes or antigens, have demonstrated higher efficacy when compared with ADC therapy alone (*15, 16*)(*17*), the two therapeutic agents might not reach the same cancer cell nor are they in close proximity to maximize endocytosis of the therapeutic ADC payload (*1, 6, 18*). In the context of bispecific antibodies, one bispecific (*19*) has attained regulatory approval to date for the treatment of solid tumors, with suboptimal pharmacokinetics, distribution, and significantly higher costs limiting translation compared to conventional antibody therapies (*14*). Moreover, the existing technology lacks optimization for the production of bispecific ADCs (*20*), and as of now, there are no clinically approved bispecific antibodies successfully conjugated with cytotoxic drugs in a manner resembling ADCs used in the clinics (*21*).

To address the challenges associated with single-antigen targeting and the complexities of bispecific antibodies, we developed an “antibody-ADC click” strategy leveraging the potential of *in vivo* click chemistry to allow for dual-targeting in tumors. This technology involves sequential administration of FDA-approved mAbs, followed by the *in vivo* click through the precise biorthogonal click chemistry — recognized by the 2022 Nobel Prize (*22–27*) — in a living organism. Notably, bioorthogonal click chemistry creates unique opportunities to allow the development of therapies targeting distinct antigens using selective reactions *in vivo* (*22, 23*). In this context, recent studies highlight clinical promise in utilizing tetrazine (Tz)-transcyclooctene (TCO) cycloaddition reaction for pre-targeted payload delivery (*28*).

Our study advances the use of combinatorial approaches with mAbs bearing Tz and TCO click moieties by incorporating groundbreaking innovations in bioorthogonal chemistry. This approach allows for sequential dual receptor targeting within tumors, surpassing the targeting ability of conventional ADCs. Our findings establish *in vivo* biorthogonal clicking of mAbs and ADCs, targeting distinct epitopes or antigens, as a promising approach for treating tumors non-responsive to or resistant to conventional ADC therapies. “Antibody-ADC click” reported here lays the foundation for further integration into distinct antibody and/or ADC or theranostic combinations offering multiple avenues for cancer therapy and other diseases.

## RESULTS

### Optimization of Bioorthogonal Clicking of FDA-approved Antibodies

ADCs currently approved by the FDA rely on targeting a single receptor at a time to deliver a cytotoxic payload to cancer cells (*5*). Here, we optimized FDA-approved mAbs and ADCs with bioorthogonal clicking moieties to enable effective sequential *in vivo* dual-targeting and drug delivery of heterogeneous cancers expressing multiple receptors (**Fig. 1a**). Our approach offers several advantages when compared with conventional ADC monotherapies or mAb combinations. First, antibody-ADC bioconjugation takes place within the living organism, enabling dual-targeting without the need for extensive bioengineering of the mAbs. Second, it allows for dual blockade of receptors enabling a more sustained tumor treatment compared to conventional ADC therapy. Third, the ADC targets not only cancer cells that express the target antigen, but it also clicks with the respective mAb pair on the cellular surface regardless of ADC-targeting receptor presence. Fourth, an antibody click approach results in a large number of receptor-antibody complexes at the surface of cancer cells, which enhances the rate of receptor-antibody internalization and subsequent degradation within lysosomes.To optimize mAbs capable of bioorthogonal clicking, we selected two model click pairs composed of mAbs targeting the human epidermal growth factor receptor 2 (HER2) and epidermal growth factor receptor (EGFR), which are receptor tyrosine kinases co-expressed in a panel of cancers (*29–31*). We optimized the click conjugation ratios of TCO- or Tz- modified mAbs. The antibodies were conjugated with the TCO or Tz click moieties (containing 4 and 5 polyethylene glycol spacers) via *N*-hydroxysuccinimidyl ester reactions (**Extended Fig. 1a**). The modification of mAbs with TCO or Tz did not affect antibody immunoreactivity (> 99%) or stability (> 90%, **Supplementary Fig. 1a,b)**. Mass spectrometry analyses of the conjugates showed 10-12 TCO/Tz click moieties per mAb (**Extended Fig. 1b,c**).

**Figure 1.**
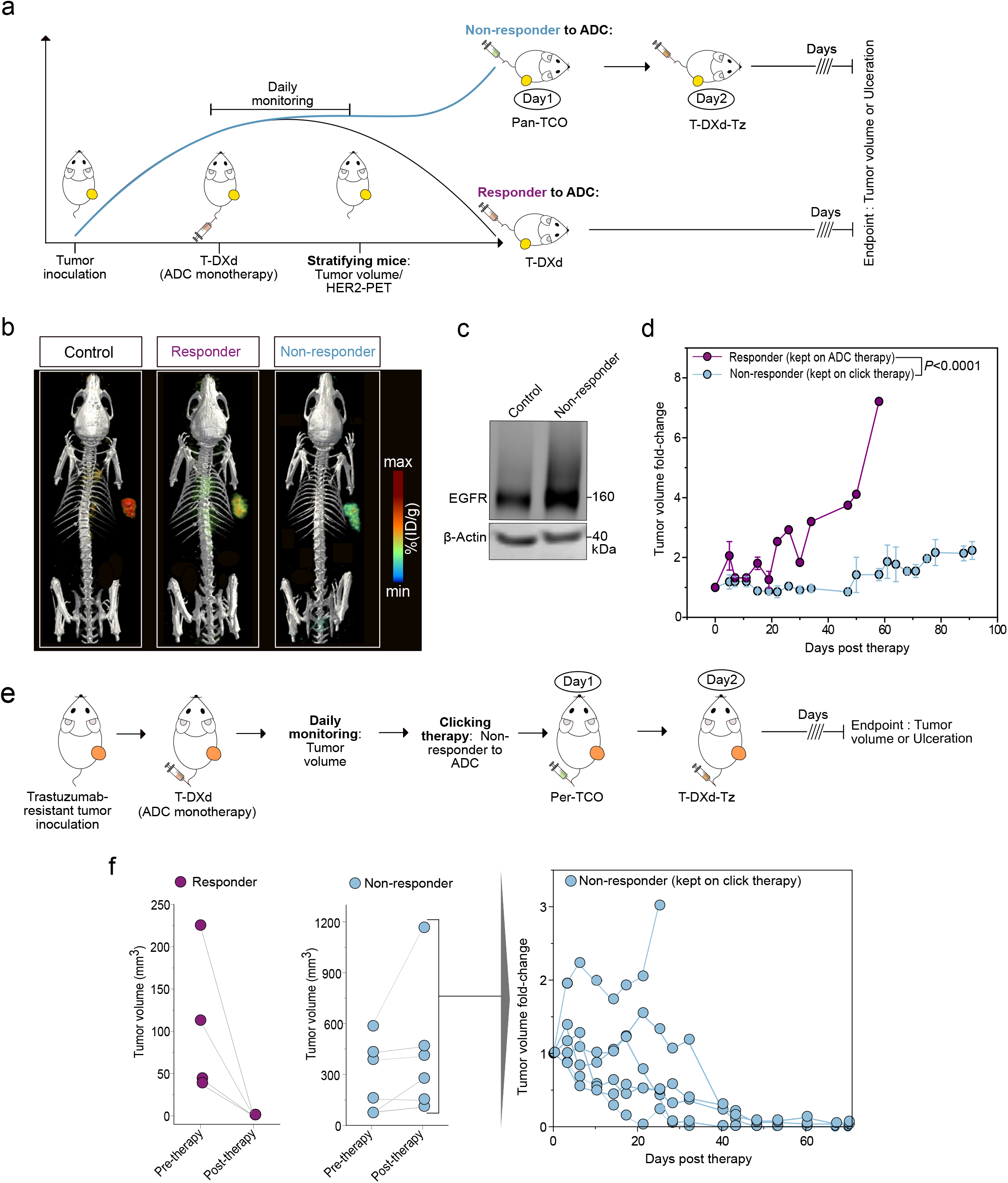
Click mAbs enable dual receptor targeting. **(a)** Schematic representation of antibodies cross-clicking on the surface of cancer cells. Clicking mAbs enables dual receptor targeting and enhances drug delivery in tumors. **(b)** SDS-PAGE analyses and quantification of the clicking antibodies at a 1:1 ratio over different incubation times (1-30 min), and different ratios (1:0.2-1:0.8) at 90 min. **(c)** Cryo-Transmission Electron Microscopy images of the no click mAbs versus clicking mAbs. Clicking mAbs were prepared by reacting antibody-TCO and antibody-Tz in a 1:1 ratio for 90 min.

In our approach, the click pair I antibody-TCO is initially administered and selectively accumulates in the tumor. Subsequently, the click pair II antibody-Tz is injected, and it selectively undergoes a click reaction with the antibody-TCO already present in the tumor (**Fig. 1a**). *In vivo* click reactions occur between antibodies that target different receptors on the cancer cell or between antibodies that target distinct regions of the same target. Being this a sequential step *in vivo*, it allows the antibody-Tz (click pair II) to still interact with the first antibody-TCO (click pair I)-target complex, regardless of antigen presence. To monitor the bioorthogonal click reactions between antibody-TCO (click pair I) and antibody-Tz (click pair II), we performed SDS-PAGE analyses (**Fig. 1b**, **Supplementary Fig. 1c**). The Tz-modified mAbs were fluorescently labeled with indocyanine green (ICG) dye to enable visualization of the click reaction. We performed click reactions at varying times (0-90 min). In other assays, we maintained a constant amount of antibody-Tz labeled with ICG in the presence of varying quantities of antibody-TCO. The click reaction between antibody-TCO and antibody-Tz occurred between 30-90 minutes when the antibodies were reacted at equimolar ratios between 1:1 and 1:0.8 of antibody-TCO: antibody-Tz (**Fig. 1b**, **Supplementary Fig. 1c**). Control experiments were conducted by adding a 15-fold excess of unreacted TCO or Tz click moieties to quench the click reaction (**Supplementary Fig. 1d)**.

Next, to validate that antibody clicking results in antibody dimerization, we performed transmission electron microscopy (TEM) imaging of the pre-clicked mAbs at a 1:1 ratio for 90 min. The TEM images demonstrated the formation of mAb dimers and clusters upon click reaction between Panitumumab-TCO (Pan-TCO, anti-EGFR mAb) and Trastuzumab-Tz (Trast-Tz, anti-HER2 mAb) (**Fig. 1c, Supplementary Fig. 1e**). Overall, these results validate bioorthogonal click of TCO- and Tz-modified mAbs, demonstrating formation of covalently linked antibody complexes.

### Bioorthogonal Click Augments Antibody Cellular Retention

Previous studies have shown that antibody combinations result in a large number of antibody-receptor complexes at the cell surface of cancer cells, which enhances the rate of antibody-receptor internalization and degradation in lysosomes (*32*). To investigate whether antibody click enhances mAb internalization, we performed immunofluorescence (IF) studies comparing a panel of HER2-targeting (trastuzumab or pertuzumab) and EGFR-targeting (panitumumab) click mAbs pairs. Control studies included no click mAbs pairs (**Fig. 2a**). IF analyses of HER2-positive and EGFR-low (HER2^+^/EGFR-low) NCIN87 cancer cells (**Supplementary Fig. 2a)** demonstrated higher intensity (*P* < 0.0001) and colocalization (∼3.5-fold; *P* < 0.0001) of the clicked anti-HER2/anti-EGFR mAbs compared to the no click mAbs pair (**Fig. 2b**). HER2^+^/EGFR^-^ BT474 cancer cells resistant to trastuzumab or CT26 cancer cells stably expressing human HER2 (CT26-hHER2) demonstrated higher uptake (*P* < 0.0001) of the anti-HER2 clicked mAbs pair targeting distinct epitopes of the HER2 receptor (pertuzumab-TCO plus trastuzumab-Tz) compared to no click mAbs (**Extended Fig. 2a**). Control studies in the CT26 mouse cancer cell line that does not express human HER2 or human EGFR demonstrated low and comparable uptake in both click and no click conditions (**Extended Fig. 2a**). Additional control studies using non-specific IgG-Tz and TCO-conjugated anti-HER2 or anti-EGFR antibodies demonstrated substantial binding of IgG-Tz to cells where anti-EGFR or anti-HER2 antibody-TCO were attached, in contrast to the no-click conditions (**Supplementary Fig. 2b)**. These results in HER2^+^ and HER2^+^/EGFR-low cancer cells demonstrate that dual engagement of two antibodies targeting distinct receptors or different epitopes of the same receptor *via* bioorthogonal click improves mAb uptake when compared to conventional single-receptor binding. Furthermore, our studies demonstrate that the antibody-Tz pair effectively engages in click chemistry with the antibody-TCO bound to the cancer cells, irrespective of the expression level of the target antigen for antibody-Tz.

**Figure 2.**
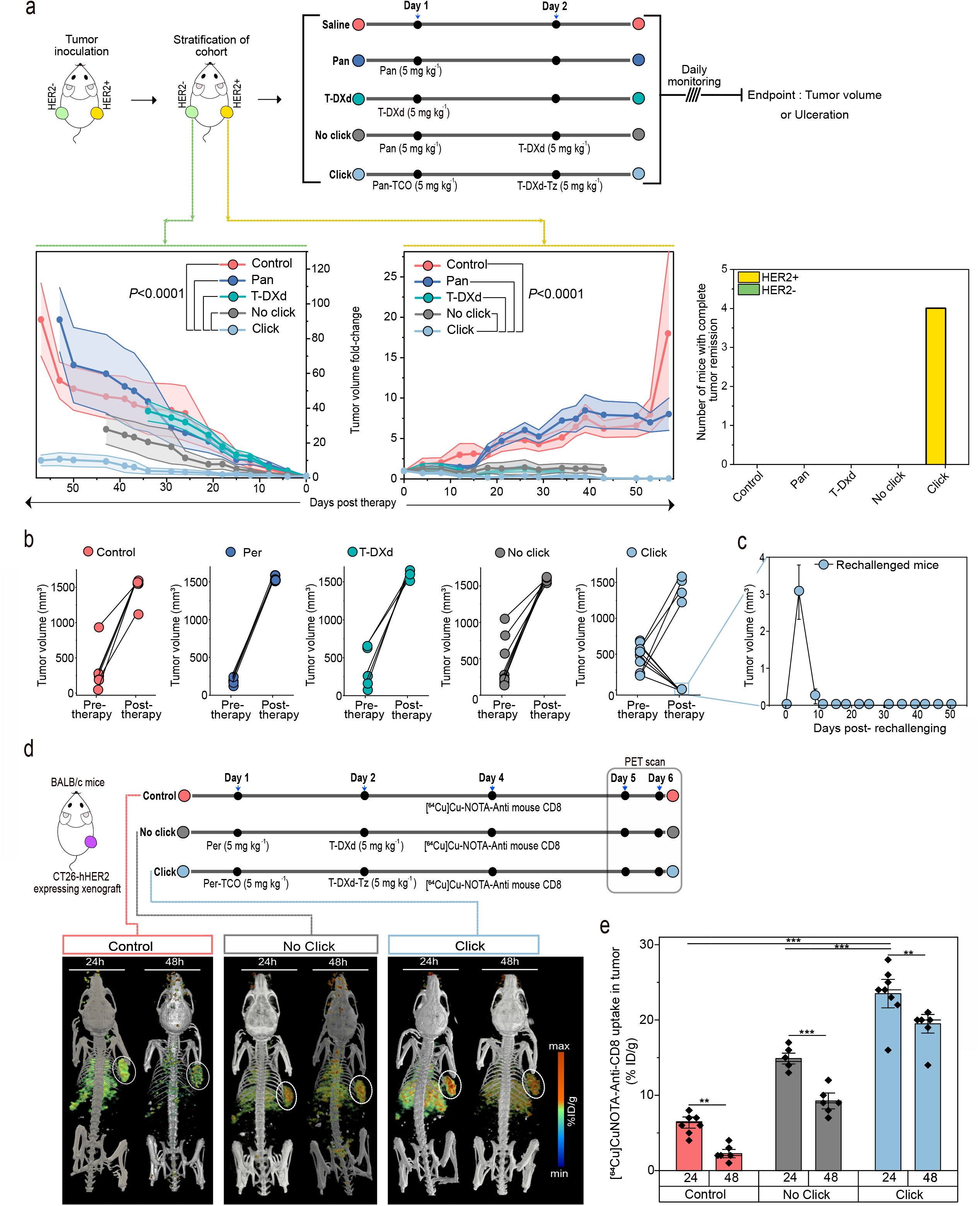
Click mAbs enhance drug delivery in cancer cells. **(a)** Schematic representation of mAbs conjugated with TCO, Tz, Alexa-488, Alexa-594, or pHrodo. **(b)** Immunofluorescence analyses of the click mAbs *versus* no click mAbs. The fluorescence intensity was obtained by quantitative analyses of fluorescence in HER2^+^ NCIN87 cancer cells cultured on 96 well plates. Quantitative analyses of the co-localized vesicles in the no click and click groups (n=10). **(c)** Fluorescence microscopy images of the no click and click groups with fluorescently labeled mAbs and LAMP1 in HER2^+^ NCIN87 cancer cells. **(d)** Fluorescence microscopy images of the no click and click groups with mAbs labeled with the pH-sensitive dye pHrodo which fluoresces upon internalization. **(e)** Quantitative analyses of the T-DM1 ADC internalization in the HER2^+^ NCIN87 cancer cells treated with no click and click mAbs at 48 h timepoint using a commercially available ELISA kit.

In EGFR^+^ A431 cancer cells that do not express HER2 receptors (HER2^-^/EGFR^+^, **Supplementary Fig. 2a)**, the click conjugation of anti-HER2-Tz mAbs with anti-EGFR-TCO mAbs allowed binding of anti-HER2 mAbs (**Extended Fig. 2a**). These results suggest that the anti-EGFR-TCO facilitates mAbs binding via biorthogonal click with anti-HER2 antibody-Tz in A431 cells. Altogether, these findings suggest a potential to extend the targeting of anti-HER2 mAbs therapy beyond cells that only express HER2, as long as it is clicked to a partnering antibody that recognizes another antigen on the cells of interest.

To further assess the endocytic trafficking of antibody, click, we utilized lysosome-associated membrane protein 1 (LAMP1) immunostaining to visualize lysosomes and late endosomes. IF analyses in NCIN87 HER2^+^/EGFR-low cancer cells revealed higher colocalization of the fluorescently tagged clicking mAbs with LAMP1-positive vesicles compared to non-clicking mAbs controls (**Fig. 2c**). To visualize antibody endocytic trafficking, we used pHrodo, a pH-sensitive dye that is non-fluorescent initially but becomes fluorescent upon internalization into acidic endosomes. We conjugated both the click and no-click mAbs pairs with pHrodo (**Fig. 2a**) and monitored internalization by IF. Clicked mAbs conjugates showed significantly higher (4-fold) pHrodo intensity versus non-clicked mAbs, indicating improved internalization and trafficking to endolysosomes (**Fig. 2d**, **Extended Fig. 2b**). Additional studies using anti-EGFR mAbs and the anti-HER2 ADC trastuzumab emtansine (T-DM1) (*33*) demonstrated an increase (3.3-fold) in DM1 drug delivery in click conditions when compared with no click (**Fig. 2e**). Overall, these data provide additional evidence that bioorthogonal click of mAbs targeting two different receptors or distinct epitopes of the same receptor enhances intracellular drug delivery compared to traditional single receptor targeting approaches.Antibody-PET Reveals Dual-Receptor Tumor Targeting by Clicking mAbs

The combination of radiolabeled antibodies with positron emission tomography (PET) allows visualization of mAbs tumor uptake and distribution in the whole body (*34*). In this study, click versus no click mAbs targeting HER2 and EGFR were radiolabeled with the positron emitters zirconium-89 (^89^Zr) or copper-64 (^64^Cu) for PET imaging followed by tissue biodistribution. As outlined in the scheme of **Fig. 3a**, mice bearing HER2^+^/EGFR-low NCIN87 tumors (**Supplementary Fig. 2a**) were administered an intravenous injection of the FDA-approved anti-EGFR antibody Pan-TCO on day 1, followed by injection of the radiolabeled anti-HER2 antibody [^89^Zr]Zr-DFO-Trast-Tz (200 μCi) on day 2. Immuno-PET and biodistribution studies at 48 h after injection of the radiolabeled immunoconjugate (**Fig. 3b,c**) revealed that the biodistribution patterns of the clicking mAb pairs exhibited tumor uptake (26.4 ± 7.5 %ID/g) and liver clearance (9.9 ± 0.66 %ID/g). These profiles were consistent with the conventional PET and biodistribution of a no click ^89^Zr-labeled mAb.

**Figure 3.**
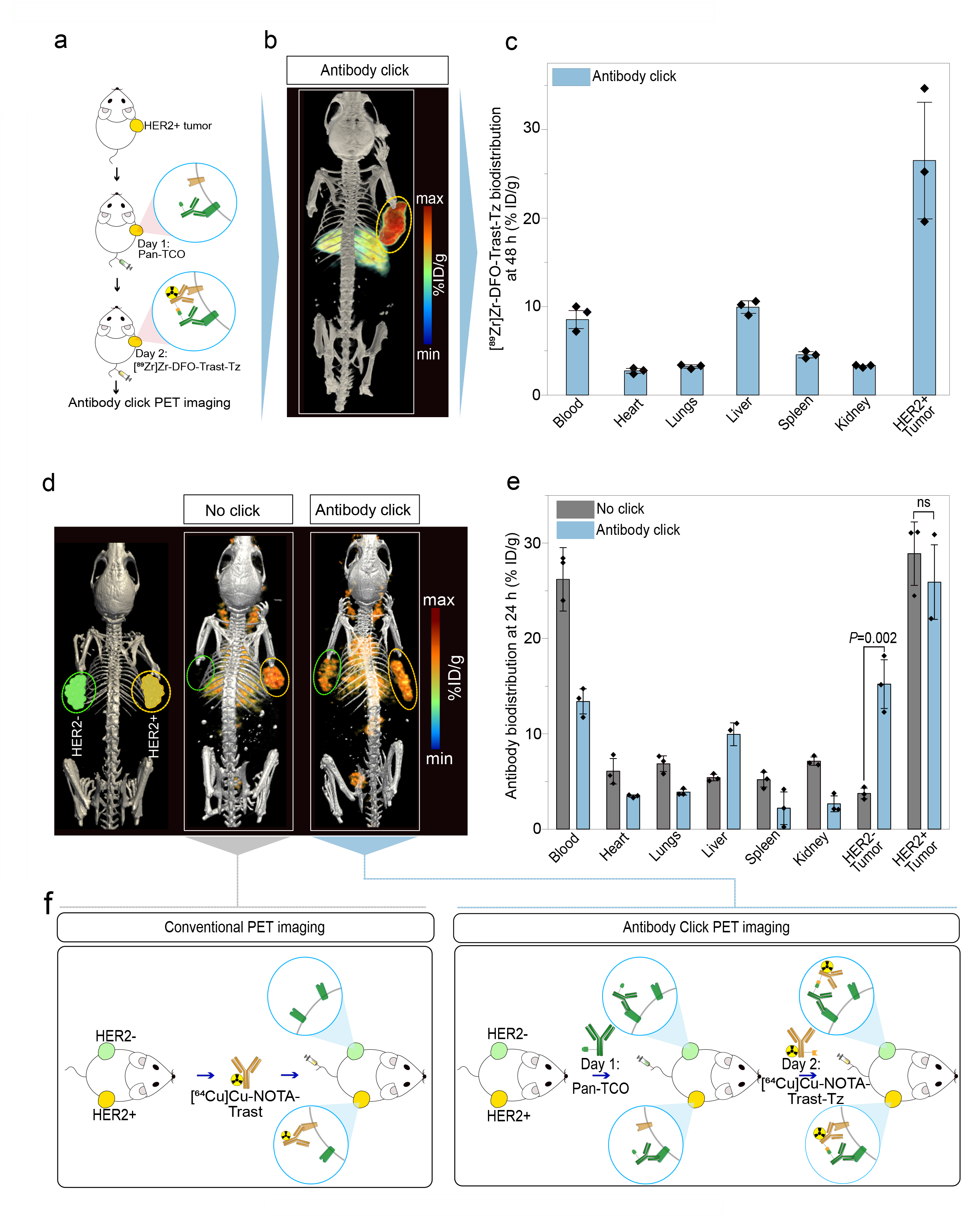
Click mAbs enable antibody accumulation in HER2^+^ and HER2^-^ tumors. **(a)** Schematic representation of the click PET imaging in the NCIN87 gastric cancer model (HER2^+^/EGFR-low). The first pair of the clicking antibodies *i.e*., Pan-TCO was injected via the tail vein 24 h prior to the [^89^Zr]Zr-DFO-Trast-Tz and PET images were acquired at 48 h post injection. **(b)** Representative PET maximum intensity projection (MIPs) image of the mice bearing the NCIN87 (HER2^+^/EGFR-low) tumor injected with the clicking antibodies and imaged at 48 h timepoint (n=3). **(c)** Biodistribution data at 48 h post-injection of clicking mAbs. The first pair of the clicking antibodies *i.e*., Pan-TCO (50 µg) was injected via the tail vein 24 h prior to the [^89^Zr]Zr-DFO-Trast-Tz (50 µg, ∼7.4 Mbq). Biodistribution was performed at 48 h post injection. Bars, n=3 mice per group, mean ± S.E.M. %ID g^−1^, percentage of injected dose per gram. **(d-e)** Representative PET MIPs image of a bilateral tumor model: NCIN87 tumors (HER2^+^/EGFR-low) on the right flank and A431 tumors (HER2^-^/EGFR^+^) tumors on the left flank. The first pair of click antibodies *i.e.,* Pan-TCO or Trast-TCO was injected *via* the tail vein 24 h prior to the injection of [^64^Cu]Cu-NOTA-Trast-Tz (50 µg, ∼7.4 Mbq). All the mice were imaged at 24 h timepoint followed by a biodistribution study. Bars, n=3 mice per group, mean ± S.E.M. %ID g^−1^, percentage of injected dose per gram. The statically significant differences between experimental groups were performed using analysis of variance (ANOVA) followed by Student’s T-test. **(f)** Schematic representation of single antibody targeting versus dual antibody click targeting. Single receptor targeting allows imaging of one receptor at a time whereas antibody-click PET allows imaging of different receptors expressed in the same tumor or in tumors inoculated in different parts of the living subject.

In addition to determining Pan-TCO/Trast-Tz in HER2^+^/EGFR-low NCIN87 cancer cells, we also evaluated the clicking of a pair of mAbs that target distinct epitopes of the HER2 receptor. We used Trast and pertuzumab (Per) that bind domains IV and II of HER2 receptor, respectively (**Extended Fig. 3a**) (*35, 36*). Antibody-PET and biodistribution studies using Trast-TCO or Per-TCO in combination with [^89^Zr]Zr-DFO-Trast-Tz demonstrated similar tumor accumulation compared to [^89^Zr]Zr-DFO-Trast-Tz alone (**Extended Fig. 3b)**. The Trast-TCO or Per-TCO binding did not result in the blocking of the second click pair *i.e.,* Trast-Tz (**Extended Fig. 3c-d)**. On the other hand, tumor uptake of mAbs binding different antigens (EGFR and HER2) Pan-TCO/Trast-Tz was ∼1.5-fold higher when compared with Trast alone (**Extended Fig. 3d)**, and further studies were conducted with this mAbs click pair.

In our pilot studies using immunocompromised mice, the control cohort treated with IgG-TCO as the click pair I demonstrated low uptake in the tumor (**Supplementary Fig. 3**). Additionally, liver uptake of click mAbs was ∼3-fold higher when compared with [^89^Zr]Zr-DFO-Trast-Tz no click (**Supplementary Fig. 3**). Antibody click in blood circulation likely enhances the interaction of the Fc domain of mAbs with Fc receptors in the liver of immunocompromised mice (*37*). Additional studies demonstrated that pre-injection of a 15-fold excess of cold IgG significantly lowered liver accumulation (**Extended Fig. 3**). Interestingly, we observed lower liver uptake of click antibodies in immunocompetent mice (8.9±1.5 %ID/g) when compared with immunocompromised mice (4.5±2.4 %ID/g), as immunocompetent mice have circulating IgG (*37, 38*) (**Supplementary Fig. 4)**.

### Antibody click enables uptake of both HER2^+^ and HER2^-^ tumors

We next sought to investigate the potential of the antibody click in an inter-tumor heterogeneous model. For this, we used a bilateral model where HER2^+^/EGFR-low (NCIN87) and HER2-/EGFR+ (A431, **Supplementary Fig. 2a**) tumors were xenografted on the right and left shoulder of *nu/nu* mice, respectively (**Fig. 3d**). This model enabled the evaluation of the uptake of the mAb click pair in HER2^+^ and HER2^-^ tumors simultaneously in the same animal. An intravenous injection of Pan-TCO was administered on day 1, followed by an injection of the anti-HER2 [^64^Cu]Cu-NOTA-Trast-Tz on day 2. The combination of Pan-TCO plus antibody [^64^Cu]Cu-NOTA-Trast (no click) was used as a control. As observed in **Fig. 3d,e**, [^64^Cu]Cu-NOTA-Trast (no click) accumulated in the HER2^+^ tumor as expected (28.9±3.8 %ID/g) with very low uptake in the HER2^-^ tumor (3.7±0.6 %ID/g). In the click group, we observed antibody uptake in both HER2^+^ (25.9±3.8 %ID/g) and HER2^-^ tumors (15.2±2.9 %ID/g). Because A431 tumors do not express HER2, the HER2-targeting mAb uptake in the HER2^-^ tumors was a result of [^64^Cu]Cu-NOTA-Trast-Tz click with Pan-TCO bound to EGFR receptors in these tumors; *i.e.* Pan accumulates in A431 tumors at 27.9±5.7 %ID/g (**Extended Fig. 4**). To further validate that a TCO/Tz approach of antibody click occurs *in vivo*, we performed additional studies using Pan-TCO or Trast-TCO followed by bioorthogonal click with control [^64^Cu]Cu-NOTA-IgG-Tz (**Extended Fig. 4**). As predicted, antibody click occurred predominantly in tumors pre-bound by the first click mAbs (EGFR-targeting panitumumab or HER2-targeting trastuzumab), validating that the TCO/Tz click reaction occurs *in vivo*. Overall, mAbs bioorthogonal click chemistry enables targeting and further imaging of both HER2^+^ and HER2^-^ tumors (**Fig. 3 e,f**).

### Antibody-ADC Click Enhances Therapeutic Efficacy

Given the enhanced tumor accumulation and internalization of antibody click when compared with the conventional no-click combinations (**Figs. 1-3**), we sought to perform therapeutic studies evaluating the efficacy of antibody-ADC click. The anti-HER2 ADC trastuzumab derutexan (T-DXd) has garnered significant recent attention due to its exceptional clinical efficacy in targeting cancer cells with potent payloads. Therefore, T-DXd was used in our therapeutic studies as the ADC-Tz click pair. First, we validated the cytotoxic potency of antibody-TCO/ADC-Tz click *in vitro* using cancer cell lines expressing varying levels of HER2 and EGFR (**Supplementary Fig. 2a)**. The 50% inhibitory concentration (IC_50_) values of T-DXd in HER2^+^ NCIN87 and HER2^-^ A431 cancer cells were 35-fold and 21-fold lower, respectively, for antibody-ADC click when compared with no click control (**Extended Fig. 5**). The results obtained in cell death assays were consistent with enhanced intracellular ADC delivery as observed in our cell studies (**Fig. 2e**).

We next validated antibody-ADC click efficacy *in vivo* using HER2^+^/EGFR-low NCIN87 tumor-bearing mice. Tumor weights at 20 days after initiating therapy were 3.7-fold and 2.8-fold higher, respectively in Pan/T-DXd (no click) or T-DXd alone (**Supplementary Fig. 5**) when compared with Pan-TCO plus T-DXd-Tz (click) at a clinically relevant dose of 10 mg kg^-1^ (*44, 45*). In subsequent therapeutic studies, we reduced T-DXd dose from 10 mg kg^-1^ to a single dose of 5 mg kg^-1^ with the aim of determining the feasibility of T-DXd dose reduction in click approaches while mitigating possible ADC-related side effects.

T-DXd shows clinical benefit in patients with HER2-high and HER2-low tumors (*39, 43*), but most of the patients will ultimately experience disease progression. Additionally, T-DXd does not accumulate in high levels in HER2^-^ cancer cells (*43*). To determine if antibody-ADC click could extend the potential benefit of an ADC to non-HER2 expressing tumors, we performed therapeutic studies in a bilateral model of HER2^+^ NCIN87 and HER2^-^ A431 cancer cells (**Fig. 4a**, **Supplementary Fig. 6**). Mice received an intravenous injection of Pan or T-DXd (5 mg kg^-1^ once) alone or in combination. The anti-tumor response in A431 and NCIN87 tumors was higher in click groups when compared with the other tested groups (*P* < 0.0001, **Supplementary Fig. 7**). T-DXd alone inhibited tumor growth in HER2^+^ tumors (*P* = 0.0757, T-DXd versus click in NCIN87) but HER2-negative tumors did not respond to T-DXd and developed ulceration (*P* < 0.0001, T-DXd versus click in A431). The combination of Pan with T-DXd (no click) slightly decreased tumor volume in A431 HER2^-^ tumors, when compared with T-DXd monotherapy, but tumor ulceration in the no-click group was observed at 34 days after therapy. Mice receiving a combination of Pan-TCO and T-DXd-Tz (click, both at 5 mg kg^-1^), showed complete remission of the HER2^+^ tumors in 80% of total treated mice. Among all the tested groups in A431 HER2^-^ tumors, the Pan-TCO/T-DXd-Tz click group exhibited the most potent inhibition of tumor growth (*P* < 0.0001, **Supplementary Fig. 8**). Additionally, the T-DXd click led to enhanced survival compared to both T-DXd alone or in combination with Pan (no click, *P* < 0.0001, **Supplementary Fig. 6)**. While we observed differences in tumor response in males versus female mice, the click-treated group demonstrated superior performance in both male and female subjects for both A431 and NCIN87 tumors (*P* < 0.0001, **Supplementary Fig. 9-11**). These results further demonstrate the potential of antibody-ADC click to augment ADC efficacy against tumors of heterogeneous expression of the targeted antigen.

**Figure 4.**
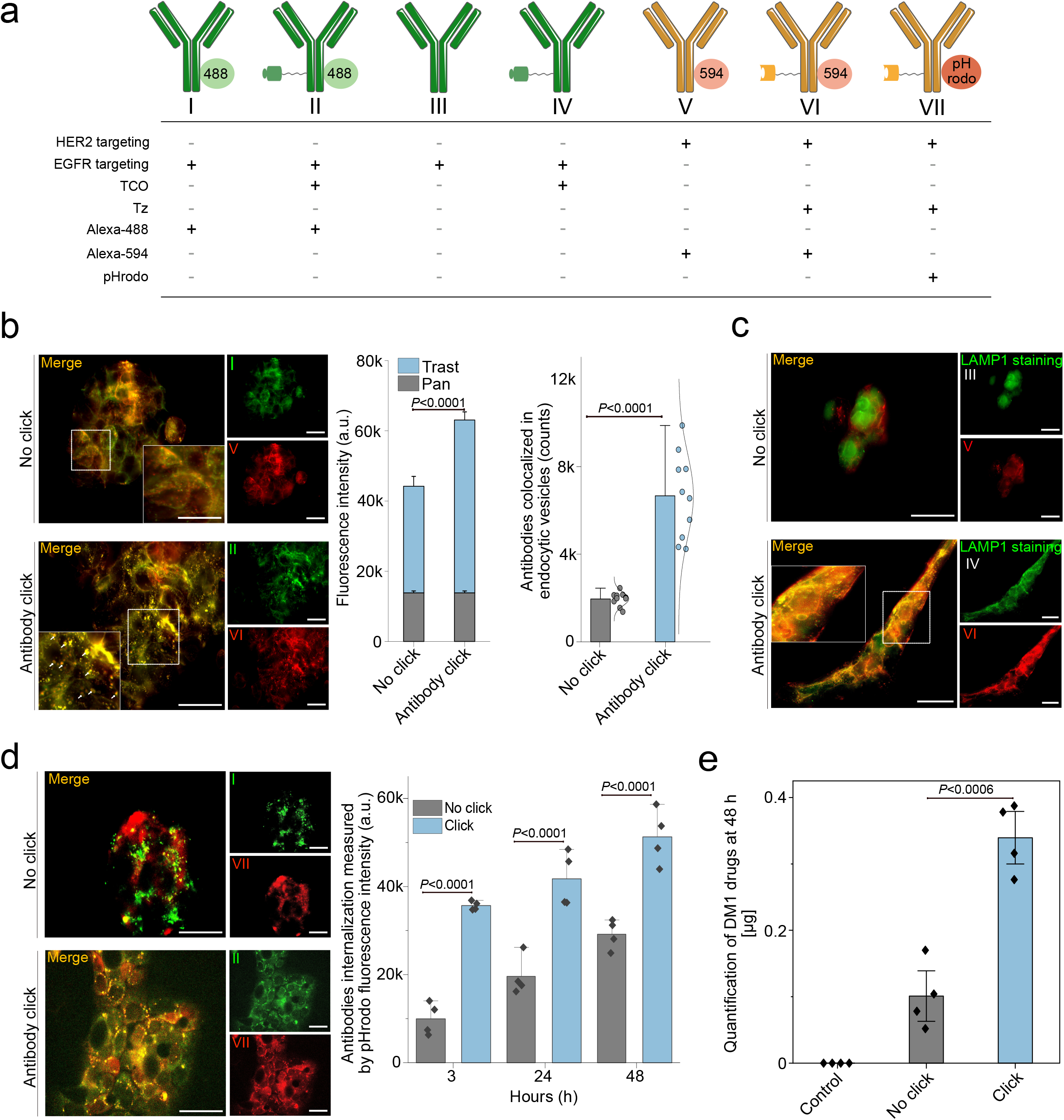
Antibody click enables therapy of HER2^+^ and HER2^-^ tumors. **(a)** Schematic of a bilateral tumor model where NCIN87 (HER2^+^) and A431 (HER2^-^) cancer cells were xenografted on the right and left shoulder of the mice, respectively. Mice bearing HER2^+^/HER2^-^ bilateral tumors were stratified into different cohorts: saline, Pan, T-DXd, no click (Pan plus T-DXd), and antibody-ADC click (Pan-TCO + T-DXd-Tz). A single dose of intravenous injection of each antibody was started at day 0. For the mice receiving the combination of drugs, the first pair of click antibodies *i.e.,* Pan or Pan-TCO (5 mg kg^-1^) was injected *via* the tail vein 24 h prior to the injection of T-DXd or T-DXd-Tz (5 mg kg^-1^). Bars, n=5 male mice per group, mean ± standard error. Therapeutic data obtained in female mice (n=5 female mice per group) is shown in **Supplementary Figure 9**. **(b)** *In vivo* therapeutic efficacy of different cohorts (*i.e.,* control saline, Per, no click Per + T-DXd and antibody-ADC click Per-TCO plus T-DXd-Tz) validated in immunocompetent mice (BALB/c) bearing CT26-hHER2 tumors. A single dose of intravenous injection of each antibody was started at day 0. For the mice receiving the combination of antibodies, the first antibody click pair *i.e.,* Per or Per-TCO (5 mg kg^-1^) was injected *via* the tail vein 24 h prior to the injection of T-DXd or T-DXd-Tz (5 mg kg^-1^). Bars, n= 10 mice per group, mean ± standard error. **(c)** Rechallenging of mice with complete tumor remission after antibody-ADC click therapy performed by xenografting the CT26-hHER2 cells on the left shoulder. Bars, n= 6 mice per group, mean ± standard error. **(d-e)** CD8-targeted PET imaging of tumor-infiltrating T cells in mice treated with the click and no click therapy. Scheme represents the mice receiving the combination of drugs: the first pair of click antibodies *i.e.,* Per or Per-TCO (5 mg kg^-1^) was injected *via* the tail vein 24 h prior to the injection of T-DXd or T-DXd-Tz (5 mg kg^-1^), whereas control received saline intravenous injections. All the mice were then injected with [^64^Cu]Cu-NOTA-anti-mouse CD8 (∼50 µg, ∼7.4 Mbq) on day 4 followed by PET imaging on days 5 and 6. The [^64^Cu]Cu-NOTA-anti-mouse CD8 tumor uptake was calculated based on the region of interest (ROI) drawing based on the PET images acquired at 24 h and 48 h timepoint. Bars, n=8 mice per group, mean ± standard error. The statistically significant differences between experimental groups were performed using analysis of variance (ANOVA) followed by Student’s *t*-test.

In addition to NCIN87 xenografts, we evaluated antibody-ADC click therapy in immunocompetent mice bearing CT26-hHER2 tumors (**Supplementary Fig. 2a)**. When mice with CT26-hHER2 were treated with the control vehicle, monotherapies 5 mg kg^-1^ anti-HER2 (Per), T-DXd, or a combination of Per plus T-DXd (no click), the mean of tumor volumes at 12 days post-therapy were >1500 mm^3^ (**Fig. 4b**). A complete response was observed in 60% (6/10) of mice who received the Per-TCO/T-DXd-Tz click treatment. No complete responses were observed in the no-click cohorts. Subsequently, the mice in the click cohort that showed complete remission of the tumors were rechallenged with CT26-hHER2 cancer cells. We did not observe tumor growth in the rechallenged mice (**Fig. 4c**), potentially indicative of enhanced immune cell infiltration. When CD8+ T cells imaging were conducted, CT26-hHER2 tumors treated with antibody click demonstrated, a ∼3.6 and 1.5-fold increase in the accumulation of CD8^+^ T cells compared to vehicle or no-click-treated groups as quantified by CD8-targeting PET imaging (**Fig. 4d,e**). Additional *in vitro* studies in cancer cells co-incubated with peripheral blood mononuclear cells from a healthy donor demonstrated an increase in cell lysis in antibody-ADC click when compared with no click (**Supplementary Fig. 12**). These results suggest that antibody-ADC click strategies using the anti-HER2 T-DXd ADC result in enhanced intracellular drug delivery, potent ADC cytotoxicity in both HER2^+^ and HER2^-^ tumors, and antitumor memory response in heterogeneous tumor models.

### Bioorthogonal Antibody-ADC Click Enables Efficacy Against Unresponsive or Drug-Resistant Tumors

Having validated augmented ADC accumulation *via* bioorthogonal antibody-ADC click, we next evaluated efficacy in cancer models unresponsive or resistant to ADC therapy. First, mice bearing HER2^+^/EGFR-low NCIN87 xenografts were administered T-DXd monotherapy. At 30 days after therapy, mice were divided into T-DXd responders (fold-change in tumor volume equal to or lower than 1) and non-responders (fold-change in tumor volume higher than 1) as shown in **Fig. 5a** and **Supplementary Fig. 13a**. HER2-targeting immuno-PET studies using [^64^Cu]Cu-NOTA-Trast demonstrated a 45% and 73% reduction in HER2 expression in T-DXd responder and non-responder tumors, respectively, as observed by immunoPET (**Fig. 5b** and **Supplementary Fig. 13b**).

**Figure 5.**
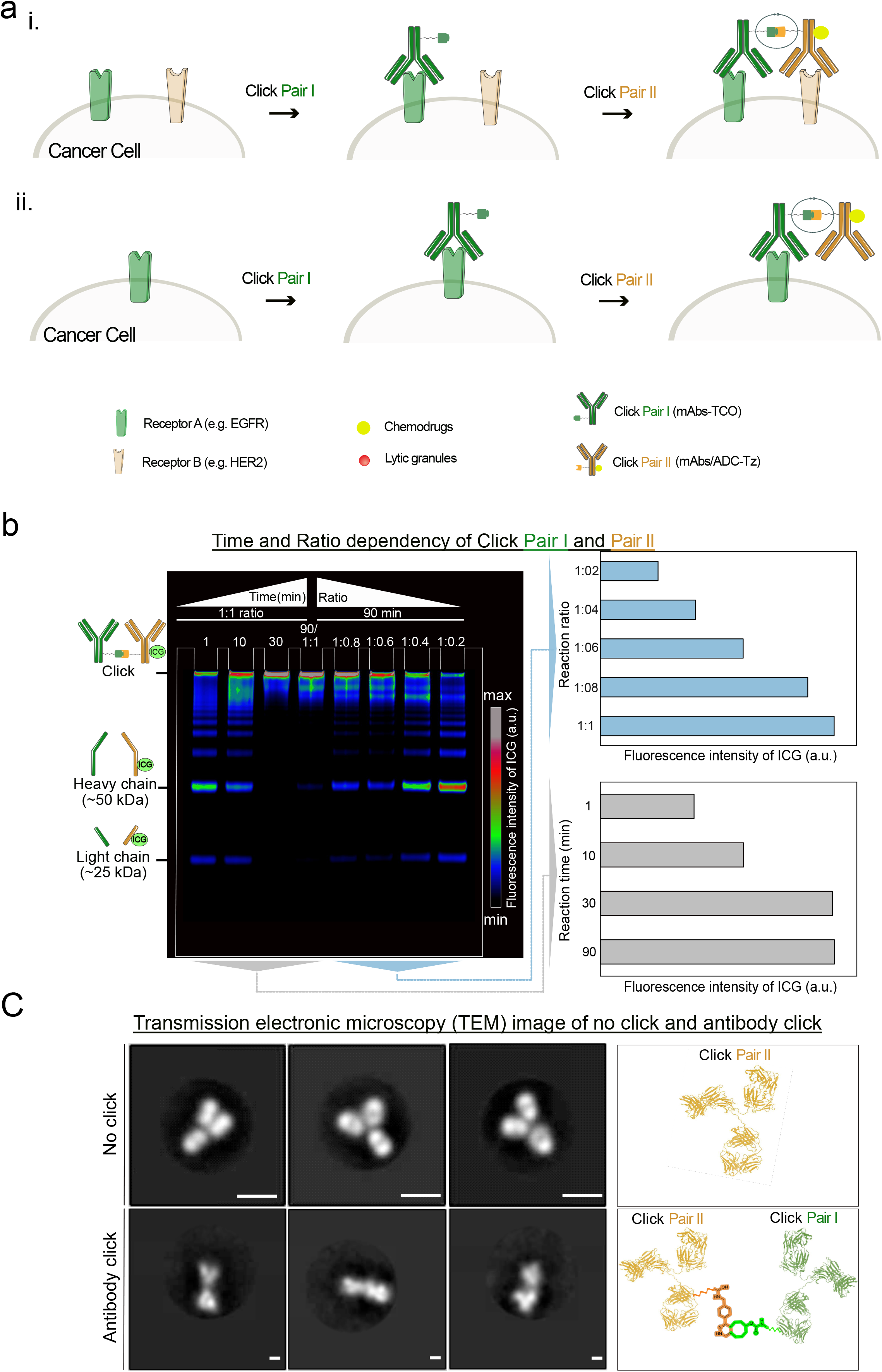
Antibody click reverses drug resistance in tumors. **(a)** Schematic of experimental treatment protocol to study the efficacy of click therapy in tumors resistant to T-DXd therapy. Mice xenografted with NCIN87 gastric cancer cells were treated with T-DXd (5 mg kg^-1^). The mice were then stratified as responders and non-responders based on tumor volume measurements. Later, the responder mice were kept on the T-DXd therapy whereas the non-responder mice were treated with the click therapy. For click therapy, the first pair of click antibodies *i.e.,* Pan-TCO (5 mg kg^-1^) was injected *via* the tail vein 24 h prior to the injection of T-DXd or T-DXd-Tz (5 mg kg^-1^). Bars, n=4-8 male mice per group, mean ± S.E.M. Therapeutic data obtained in female mice (n=5 female mice per group) is shown in **Supplementary Figure 13**. **(b)** Anti-HER2 PET imaging was performed to monitor HER2 protein levels in the responder versus non-responder tumors in male mice upon stratification on day 30. The mice were intravenously injected with [^64^Cu]Cu-NOTA-Trast-Tz (80 µg, ∼10.5-11.84 Mbq) and PET imaging was performed at 24 h timepoint (n=3). **(c)** Western blot showing EGFR upregulation in T-DXd non-responder tumors when compared with control tumors. **(d)** The responder mice were kept on the T-DXd therapy (5 mg kg^-1^) whereas the non-responder mice were kept on the click therapy. For click therapy, the first therapy of click antibodies *i.e.,* Pan-TCO (5 mg kg^-1^) was injected *via* the tail vein 24 h prior to the injection of T-DXd or T-DXd-Tz (5 mg kg^-1^). **(e,f)** Efficacy of antibody-ADC click in tumors resistant to trastuzumab therapy. The mice were xenografted with Trastuzumab-resistant BT474 cancer cells. The mice were then stratified into responders and non-responders based on tumor volume measurements. The non-responder mice were treated with a click therapy. For click therapy, the therapy of click pair *i.e.,* Per-TCO (5 mg kg^-1^) was injected *via* the tail vein 24 h prior to the injection of T-DXd or T-DXd-Tz (5 mg kg^-1^).

Prior clinical data demonstrated EGFR upregulation as a resistance mechanism to HER2-targeted therapies (*31*), and we, therefore, assessed the EGFR protein levels in tumors that exhibited non-responsiveness to T-DXd. Western blot studies showed a 2.2-fold increase in EGFR total protein levels in the non-responder T-DXd treated tumors when compared with responders (**Fig. 5c**). After stratification in ADC responders versus non-responders (*P* = 0.0092), the responders remained on T-DXd monotherapy, while the non-responder mice were switched to a combination of anti-EGFR Pan-TCO plus anti-HER2 T-DXd-Tz click therapy. Notably, antibody-ADC click therapy reduced tumor growth in previously non-responder tumors compared to mice that continued to receive ADC monotherapy (*P* < 0.0001). In cohorts treated with no click ADC, tumors transitioned from responder to non-responders (**Fig. 5d**). In this study, we observed a trend suggesting an improved therapeutic response to click therapy in male mice when compared to female mice (**Supplementary Fig. 13a-c**). However, statistical analysis revealed no significant difference in click response between males versus female mice (**Supplementary Fig. 14**).

To further confirm the potential of antibody-ADC click therapy to treat resistant tumors, we validated click efficacy in a BT474 trastuzumab-resistant tumors (**Fig. 5e**). Because BT474 trastuzumab-resistant cancer cells express HER2, we used the anti-HER2 mAb Per in combination with T-DXd as the click approach. Additionally, prior research has shown that T-DXd plus Per is superior to T-DXd alone in treating HER2-expressing tumors (NCT04784715) and others have shown T-DXd efficacy in tumors resistant to trastuzumab (*43, 46*). Mice inoculated with trastuzumab-resistant BT474 tumors were treated with T-DXd alone and stratified as responder or non-responders based on their tumor volume (**Fig. 5e**, **Supplementary Fig. 13d**). Six out of 10 mice bearing BT474-trastuzumab resistant tumors demonstrated to be non-responders to T-DXd. Non-responder mice were then switched to a combination of Per-TCO and T-DXd-Tz click. Upon switching BT474-trastuzumab resistant tumors that exhibited resistance to both trastuzumab and T-DXd, to an antibody-ADC click therapy, the antibody-ADC click pair demonstrated effective tumor suppression (*P* < 0.0001, **Fig. 5f)**. These results demonstrate that a strategy of dual targeting through antibody-ADC click effectively overcomes drug resistance, further validating the potential of this approach to enhance therapy in refractory cancers.

## DISCUSSION

While ADCs have shown promise in cancer treatment (*3–5, 15–17, 47*), they are inherently limited by their ability to target a single antigen. In this study, we introduce an antibody-ADC click approach that overcomes the challenges associated with tumoral heterogeneity in target expression, a limitation known to affect the efficacy of traditional ADCs. Antibody-ADC click demonstrates an enhancement in efficacy, surpassing what can be achieved through a combination of two distinct antibodies. An antibody click approach significantly enhances intracellular drug delivery without necessitating tumors to overexpress the target antigen. Furthermore, the antibody-ADC click approach streamlines the process of generating ADC biomolecules for receptor multi-targeting, eliminating the need for labor-intensive procedures. Leveraging bioorthogonal click chemistry, inspired by the 2022 Nobel Prize in Chemistry, we achieve exceptional reactivity, facilitating a highly controlled and efficient method for receptor multi-targeting and drug delivery.

The antibody click strategy we describe addresses several problems with the conventional ADCs: first, it enables multi-targeting to overcome target heterogeneity (**Fig. 1**) as shown in our studies using the HER2-targeting T-DXd. T-DXd, one of the most potent ADCs with extensive clinical approvals, represents a remarkable advancement in the field of targeted therapeutics (*39, 41–43*). T-DXd is effective in treating HER2-expressing tumors, and it has exhibited modest activity in tumors characterized by low HER2 expression. Additionally, most of the patients treated with T-DXd will ultimately experience disease progression and die. In this study, we show the potential of the click approach to expand the therapeutic benefits of T-DXd to tumors that do not express HER2 (**Fig. 4**), significantly increasing the population of tumors that can benefit from this potent ADC.

Another critical problem our approach aims to address is the simplification of the process for generating ADC biomolecules capable of achieving receptor multi-targeting without time-consuming or labor-intensive procedures. Presently, the landscape of bispecific antibody development offers an array of over 90 different technologies aimed at enabling receptor multi-targeting (*20*). However, a substantial fraction of these bispecific mAb face challenges in terms of development for clinical use or compatibility with ADC generation. A significant challenge in this domain is the creation of bispecific ADCs with favorable binding kinetics for the targeted receptors, the incorporation of an appropriate spacer between the antibody and the cytotoxic moiety, and the effective conjugation of cytotoxic payloads that will result in effective drug delivery. This multifaceted challenge has been the focal point of our work, where we harnessed the potential of ‘bioorthogonal’ click chemistry to develop antibody-ADC click conjugates (*48, 49*). The click chemistry used in our work was inspired in the 2022 Nobel Prize in Chemistry to Professors K. Barry Sharpless, Morten Meldal, and Carolyn Bertozzi. Notably, we harnessed the remarkable speed of the inverse electron-demand Diels–Alder (IEDDA) reaction, characterized by k_2_ values exceeding 100,000 M^−1^s^−1^. This exceptional reactivity made it the ideal choice for generating our antibody-ADC click system. In the antibody click approach, two targeting FDA-approved antibodies are introduced sequentially in a living biological system (**Fig. 3**), eliminating the need for extensive antibody modification that might alter their binding properties. This selective ligation strategy facilitates the *in vivo* combination of the two antibodies specifically at the tumor site, offering a highly controlled and efficient method for achieving receptor multi-targeting. Utilizing FDA-approved mAbs and the IEDDA ligation between Tz and TCO that is compatible with living biological systems and enables click *in vivo* (*50, 51*), we simplified the development of ADCs to effectively address target heterogeneity in tumors.

An antibody click approach to receptor multi-targeting enables synergistic interactions between the mAbs and their respective targets, resulting in increased intracellular payload delivery (**Fig. 2**). This characteristic associated with antibody click distinguishes our approach from a mixture of mAbs. Notably, when the antibody click approach is employed, the property of internalization is no longer confined to the inherent characteristics of individual antigens. In this context, an antigen that exhibits slow or modest internalization, when co-targeted alongside a rapidly internalizing antigen, can be transformed into an internalizing antigen upon engagement by the click antibody. Furthermore, a distinctive advantage of the antibody click strategy, in contrast to the use of mAbs mixtures, relates to its ability to modulate the binding affinity of each targeting arm without compromising the overall binding to the tumor. This phenomenon is facilitated by avidity and dual engagement, allowing the antibody click approach to overcome the limitations associated by the binding site barrier that is known to hinder the effectiveness of mAbs targeting (*52*).

The antibody-ADC click method provides a versatile platform for developing biomolecules capable of targeting either the same antigen or distinct receptors. The potential for future optimization of the spacer that connects the mAbs to the TCO and Tz click moieties, could enable the simultaneous targeting of co-expressed receptors, whether they are in close proximity or at a certain distance in the tumor. This adaptability offers distinct advantages compared to traditional biparatopic strategies. Two biparatopic ADCs targeting HER2 (ZW-49 and MEDI4276), as well as one directed against the MET receptor (REGN5093-M114) that simultaneously engage two epitopes on the same antigen, have been evaluated in clinical trials (*53–55*). However, bispecific ADCs that concurrently bind to two distinct antigens are predominantly in the preclinical development phase (*56*) The antibody-ADC approach discussed here is based on the use of FDA-approved antibodies, which can be readily modified with easily accessible TCO or Tz moieties. Furthermore, the click reaction takes place within the living system, highlighting its potential as a promising candidate for clinical translation.

The antibody-ADC click approach reported here has the potential to simultaneously address target heterogeneity within tumors, a characteristic that is not possible with a mixture of mAbs. Leveraging the IEDDA-based click chemistry, which has left a significant mark in chemistry applications, especially in radiochemistry and drug delivery (*28, 50, 51, 57, 58*), this methodology shows promise for translation into clinical applications. The clinical translation of *in vivo* IEDDA reactions is further supported by recent studies in large subjects including companion dogs with osteodestructive lesions (*59*). Moreover, the prospect of a first-in-human demonstration of this technology is planned in the near future (*28, 58*), reinforcing its potential for rapid clinical implementation. The choice of T-DXd in the proof-of concept studies reported here, a well-established ADC with a known toxicity profile, further enhances the future clinical translation of this work. Antibody-ADC click shows a biodistribution pattern that is similar to that of mAbs (**Fig. 3**), and we therefore expect safety similar to the already FDA-approved mAbs used in the generation of the antibody-ADC click. Acknowledging the known toxicity profiles of T-DXd payloads, notably DXd in relation to lung toxicity (*60, 61*), toxicity profiles and dosing strategies in murine models expressing human HER2 are necessary for clinical translation. The effectiveness of the ADC appears to be influenced by the gender of the mice utilized in this study. However, a thorough understanding of the underlying mechanisms will necessitate further in-depth investigation. We anticipate that the antibody click approach is well-positioned to find wide-ranging applicability across ADCs and multi-targeting, offering versatile solutions for various applications, encompassing direct cell killing and antibody-dependent cell-mediated cytotoxicity (ADCC). Notably, this cost-effective methodology, with readily applicable chemicals, serves as an ideal platform for screening multiple antibody-ADC combinations, driving innovation in novel drug discoveries.

## METHODS

### Ethical compliance

Animal studies were performed in the Washington University School of Medicine animal facility in compliance with institutional guidelines and under Institutional Animal Care and Use Committee (IACUC) approved protocols (Animal Protocol 21-0087, IBC protocol 13652, PI: Pereira).

### Cell culture

Human cancer cell lines NCIN87, A431, BT474, and CT26 were purchased from the American Type Culture Collection (ATCC). All cell lines used were mycoplasma-free and cultured at 37 °C in a humidified atmosphere at 5% CO_2_. All cell culture media were supplemented with 100 units mL^-1^ penicillin and streptomycin. Details of cell culture media for respective cell lines are detailed in **Supplementary Table 1**.

#### Generation of CT26 murine cancer cells stably expressing human HER2 (CT26-hHER2)

We used our previously reported methods to develop the CT26-hHER2 cell line(*62*). Briefly, we transduced the murine cancer cell line CT26 using 8 g mL^-1^ hexadimethrine bromide (Sigma) and the media was changed 24 h later. A puromycin selection (5 μg mL^-1^) was initiated 3 days after the transduction, and was continued for a minimum of 4 days after that. Western blot analysis confirmed the increase in human HER2 expression in the CT26 cells (**Supplementary Fig. 2a)**. *Ex vivo* biodistribution analyses using radiolabeled anti-HER2 trastuzumab confirmed the expression of hHER2 in CT26-hHER2 cells implanted in the mouse (**Supplementary Fig. 4)**.

#### Generation of BT474 cancer cells resistant to trastuzumab (BT474-trastuzumab resistant)

BT474 human breast cancer cells were made resistant to trastuzumab by continually exposing the parental BT474 cancer cells to increasing concentrations (up to 15 μg mL^-1^) of the trastuzumab antibody over a period of 9 months. The development of resistance was confirmed by viability assays.

### Antibody modification and characterization

#### Antibody modification

Antibodies — Panitumumab (Pan), Pertuzumab (Per), Trastuzumab (Trast), T-DM1 and T-DXd (**Supplementary Table 2**) were obtained from the Siteman Cancer Center pharmacy. Antibodies were conjugated at a molar ratio of 15 TCO-(PEG)_4_-NHS-ester (TCO) (*i.e.,* TCO) or Tz-(PEG)_5_-NHS-ester (Tz) (*i.e.,* Tz) per mAb. All conjugation reactions were carried out in an amine-free solution (PBS, pH 8.8-9) at 37 °C with 450 rpm agitation for 1 h. Additionally, the unconjugated chemical TCO/Tz click moieties were removed from the conjugated mAbs using a size exclusion column (PD-10; GE Healthcare). The conjugate solutions were further concentrated using amicon filters with a 50 kDa cutoff.

For immunofluorescence (IF) studies, the TCO- or Tz-conjugated mAbs were then labeled with Alexa Fluor 488 (Thermo Fisher Scientific, A2042), Alexa Fluor 594 (Thermo Fisher Scientific, A20004), pH-Rhodo (Thermo Fisher Scientific, P36600), or indocyanine green (ICG) at a molar ratio of 3 fluorophores per mAbs at 37°C in PBS at pH 8.8. The fluorescently-labeled mAb conjugates were purified via size exclusion chromatography (PD-10 column; GE Healthcare) and concentrated using 50 KDa cutoff amicon filters.

The concentration of the mAb conjugates was determined using a UV-visible spectrophotometer, Pierce 660 assay (Thermo Fisher Scientific, 22660), or BCA assay (Thermo Fisher Scientific, 23227). All mAb conjugates were stored at -80 °C protected from light until used.

#### SDS-PAGE gels

Pan-TCO and Trast-Tz-ICG were incubated at different ratios (1:1, 1:0.8, 1:0.6, 1:0.4 and 1:0.2), for 90 min at 37 °C with 450 rpm agitation. Additional experiments were conducted in 1:1 reaction ratios at different incubation times (90, 30, 10 and 1 min) at 37 °C with 450 rpm agitation. Then, the samples were mixed with loading buffer and boiled under reducing conditions (Laemmli buffer containing reducing agent, Invitrogen) for 5 min at 95 °C, and loaded onto SDS-PAGE gels (NuPage 4-12% Bis-Tris protein gels, Invitrogen). Alternatively, the antibody samples were mixed in loading buffer under non-reducing conditions.

To block the click reactions, mAb conjugated to trans-cyclooctene (antibody-TCO) or tetrazine (antibody-Tz) were pre-incubated with a 15-fold molar excess of unconjugated Tz or TCO, respectively. Pre-incubation was performed for 90 minutes at 37 °C with 450 rpm agitation. After pre-incubation, the clicking pair of mAbs (antibody-Tz or antibody-TCO) was added in a 1:1 molar ratio of conjugated mAbs. The degree of click conjugation was then compared to a positive control with no blocking (clicked mAbs) and a negative control with no click moieties (unconjugated mAbs). After SDS-PAGE, the gels were rinsed with deionized water, stained with Pierce Mini Gel Power Staining and Destaining Kit, and scanned on the Odyssey CLX.

#### MALDI spectrometry and TEM

Matrix-assisted laser desorption/ionization-time of flight (MALDI-TOF) mass spectrometry of the antibody conjugates was performed to determine the number of conjugates per mAbs at the Alberta Proteomics and Mass Spectrometry Facility at University of Alberta in Canada. TEM of the unconjugated and conjugated mAbs was performed at the Washington University Center for Cellular Imaging (WUCCI). Briefly, 10 μL of no-click Trast or click Trast-TCO/Trast-Tz samples (5 μg/mL in mQ water) were absorbed for 60 seconds onto carbon-coated 200-mesh copper grids (01840-F, Ted Pella), which had been glow-discharged for 30 seconds in a Solarus 950 plasma cleaner (Gatan). After sample absorption, grids were washed 5 times with ultrapure water and stained for 2 minutes with freshly prepared 0.75% uranyl formate. Excess stain was blotted off using filter paper (Whatman No.2, Fisher Scientific) before air drying. Grids were imaged with a JEOL JEM-1400Plus TEM (JEOL USA Inc) at an operating voltage of 120 kV with a NanoSprint15-MkII 16-megapixel sCMOS camera (Advanced Microscopy Techniques). For 2D classification analysis, images were collected at a nominal magnification of 30kx, corresponding to a pixel size of a 3.54 Å. Data processing was done with Relion 3.1 (PMID: 30412051). Briefly, particles were picked using Laplacian-of-Gaussian blob detection and then extracted with box sizes of 128 or 250 pixels for the ‘no-click’ and ‘click’ samples, respectively. Particles underwent multiple rounds of 2D classification, and the clearest classes were selected for display.

### Immunofluorescence

Cancer cells (1 million cells) grown on coverslips of 1.5 mm thickness (ThermoFisher Scientific, 12-542B) were first incubated with 100 nM of Alexa Fluor 488 tagged antibody-TCO in PBS containing 1% bovine serum albumin (BSA) for 30 min at 4 °. Next, cells were washed to remove unbound mAbs and incubated with 100 nM of antibody-Tz tagged with Alexa-Fluor 594 or pHrodo (Excitation: 560 nm and Emission: 587 nm) for 3, 24 or 48 h at 37 °C in 5% CO_2_. Control experiments included unconjugated mAbs labeled with Alexa Fluor 488 (Excitation: 490 nm and Emission: 525) or Alexa Fluor 594 (Excitation: 590 nm and Emission: 620 nm).

Experiments of LAMP-1 staining were performed after cells fixation with 4% paraformaldehyde (PFA) and permeabilization with PBS containing 1% Triton X-100, followed by incubation with a rabbit anti-LAMP-1 primary mAb (ab24170, Abcam) and secondary goat anti-rabbit IgG fluorescently labeled with Alexa Fluor 488 (A-11008, ThermoFisher Scientific). Fluorescence images were acquired using a 60X oil immersion objective on the EVOS M5000 Imaging microscope.

### pHrodo-internalization assays

Cells were plated in a 96-well plate (10,000 cells/well). Twenty-four hours after plating, cells were incubated with antibody-TCO-Alexa Flour 488 or mAbs-Alexa Flour 488 (100 nM) in 1% BSA in PBS for 30 min at 37 °C, followed by washing using ice-cold PBS. Later, cells were incubated with antibody-Alexa Flour 594, antibody-Tz -Alexa Flour 594, antibody-pHrodo or antibody-Tz-pHrodo (100 nM) at 37 °C and 5% CO_2_ for 3 h, 24 h and 48 h. At the time of fluorescence measurement, the cells were washed 5 times using ice-cold PBS and fluorescence measurements were recorded using the Synergy H1 microplate reader (Biotek).

### Cell viability assays

Cancer cells (20,000 cells/well: NCIN87, A431, CT26-hHER2, CT26, BT474 cells) were plated on a 96-well plate for 24 h. In no click conditions, the cells were incubated on ice with Pan, Trast or Per alone during 30 min. Next, cancer cells were washed 3 times using ice-cold PBS. Later, cancer cells were treated with T-DXd during 48 h. In antibody-ADC click treatment groups, the cancer cells were treated with TCO modified mAbs (Pan-TCO, Per-TCO, or Trast-TCO) for 30 min on ice. After washing cells during 3 times using ice-cold PBS, the cells were treated during 48 h with T-DXd drugs modified with Tz (T-DXd-Tz). Antibody concentrations were tested at 100, 10, 1, 0.1, 0.01, and 0.001 nM and at a molar ratio of 1:1 in all click and no click conditions. After 48 h of incubation with the no click or click pairs, a media change was performed, followed by the addition of 20 μL of a mixture of WST-8 (1.5 mg mL^−1^, Cayman Chemical) and 1-methoxy-5-methylphenazinium methylsulfate (100 μM, Cayman Chemical) per manufacturer’s instructions. Next, the plates were incubated for 2 h in a CO_2_ incubator with gentle agitation, and absorbance in each well was measured at 460 nm.

### ELISA assays

#### Antibody binding to human HER2 protein

HER2 protein (100 µL) at a concentration of 250 ng/mL was coated overnight on 96-well plates at 4°C. The solution was subsequently discarded, and the plates were blocked with 1% bovine BSA blocking buffer for 1 hour at 37°C. The mAbs, antibody-TCO, or antibody-Tz were then incubated overnight at 4°C. After washing four times with wash buffer, 100 μL of an anti-IgG HRP-conjugate was added to each well and incubated for 1 hour at room temperature. The plates were washed four times again, then 100 μL of 3,3’,5,5’-Tetramethylbenzidine substrate was added to each well and incubated at 37°C for 30 minutes. The reaction was stopped by the addition of stop solution (Human ErbB2 ELISA Kit, Invitrogen) and the absorbance was measured at 450 nm.

#### In vitro quantification of ADC payload intracellular delivery

Cells were plated on six-well plates (2 million cells/well). After 24 h, the cells were treated with no click or click pair of mAbs. The mAbs or antibody-TCO (100 nM) was incubated on ice for 30 min followed by washing using ice-cold PBS. Later, the T-DM1 or T-DM1-Tz (100 nM) were added to the cells on ice for 30 min under shaking conditions followed by incubation at 37 °C and 5% CO_2_ for 48 h. Samples were prepared for the DM1 ELISA assay as instructed by the manufacturer (Eagle Biosciences, KTR-756).

### *In vitro* analysis of antibody-dependent cellular cytotoxicity (ADCC) experiments

Cells were plated on 96 well plates (20,000 cells/well). After 24 h, the cells were treated with no click or click pair of mAbs. The first mAbs or antibody-TCO (100 nM) was incubated on ice for 30 min followed by washing using ice-cold PBS. Then Trast or Trast-Tz (100 nM) were added to the cells on ice for 30 min under shaking conditions. Later, the PBMCs (one million PBMCs/well) were added to achieve a ratio of 1:50 (*i.e.,* cancer cells: PBMCs) followed by incubation at 37 °C and 5% CO_2_ for 48 h. For IgG blocking cohorts, half a million of PBMCs were pre-incubated with 0.5 mg of IgG (Human isotope) for 1 h on ice, before culturing the PBMCs with cancer cells. ADCC in no click or click mAbs conditions was evaluated using the Cytotoxicity Detection Kit (LDH, Roche) as instructed by the manufacturer.

### Western blot analyses

Western blot of tumor and cell lysates were performed using our previously reported methods(*62, 63*). Primary antibodies used in this study included rabbit anti-HER2, 1:800 (ab131490; Abcam), rabbit anti-EGFR, 1:1,000 (ab52894; Abcam), and mouse anti-β-actin, 1:10,000 (A1978; Sigma). After incubation with the primary mAbs, membranes were washed with tris-buffered saline containing Tween-20 buffer for three times with gentle agitation and incubated with secondary antibodies (anti-mouse goat IgG conjugated with AlexaFluor Plus 800 or anti-rabbit goat IgG conjugated with AlexaFluor 680, Invitrogen) for 1 h at room temperature. Membranes were washed and scanned using the Odyssey CLX imaging system (LI-COR Biosciences).

### Radiolabeling and binding assays

#### [^89^Zr]Zr-DFO-Antibody-Tz

^89^Zr-oxalate used in the studies was purchased from the WUSTL Isotope Production Team. Antibodies were first conjugated with the chelator *p*-isothiocyanatobenzyl-desferrioxamine (DFO-Bz-NCS; Macrocyclics, Inc) at a molar ratio of 5 DFO per antibody at a pH of 8.8 (37 °C, 450 rpm agitation). The unconjugated DFO was removed from the antibody-DFO conjugates using a size exclusion column (PD-10; GE Healthcare) and the antibody solutions were concentrated using amicon filters with a 50 kDa cutoff filter. The purified antibody-DFO was then reacted with Tz at a molar ratio of 15 tetrazine-(PEG)_5_-NHS per mAb at a pH 8.8 (37 °C, 450 rpm agitation). Subsequently, the mAbs-DFO-Tz was purified using the size exclusion PD10 column in the chelex PBS, followed by amicon cleaning. After confirming mAb conjugation and clicking efficacy by SDS-PAGE and MALDI, mAbs-DFO-Tz was labeled with zirconium-89 at a pH 7.4 for 60 min at 37 °C. [^89^Zr]Zr-DFO-antibody-Tz used in the study had a radiochemical yield and purity above ∼95% respectively.

#### [^64^Cu]Cu-NOTA-Antibody-Tz

We purchased Copper-64 from the WUSTL Isotope Production Team. First, mAbs were buffer exchanged and concentrated in 0.1 M 4-(2-hydroxyethyl)-1-piperazineethanesulfonic acid (HEPES) buffer (pH 8.5). The purified mAbs was then reacted with a 20-fold molar excess of 2-S-(4-Isothiocyanatobenzyl)-1,4,7-triazacyclononane-1,4,7-triacetic acid (*p*-SCN-Bn-NOTA, Macrocyclics) overnight at 4 °C with gentle agitation. Afterwards, the NOTA modified mAbs were purified and buffer exchanged using PD10 columns in PBS (pH 7.4), followed by amicon filtration with a 50 kDa cutoff filter. mAbs-NOTA was then reacted with tetrazine-(PEG)_5_-NHS ester at a molar ratio of 15 per mAbs for 1 h at 37 °C with 450 rpm agitation. Additionally, the unconjugated NOTA was removed using a size exclusion column (PD-10; GE Healthcare) and the antibody solutions were concentrated in ammonium acetate buffer (pH 5.5) using amicon filters with a 50 kDa cutoff. The efficacy of mAbs to click or the presence of any aggregation was validated by performing SDS-PAGE followed by Coomassie staining. Later, mAbs-NOTA-Tz was reacted with ^64^Cu for 1 h at 37 °C with 450 rpm agitation. A mixture of 50 mM ethylenediaminetetraacetic acid (EDTA) and methanol was used in iTLC to determine a radiochemical yield of ∼95% and radiochemical purity of ∼98%, respectively.

#### Total binding assays

For total binding assays, 1 million cancer cells, were incubated in PBS containing 1% w/v bovine serum albumin (BSA, Sigma Aldrich) and 0.1% w/v sodium azide (NaN_3_, Sigma Aldrich) in the presence of 2.5 μg of Pan-TCO, Per-TCO, or Trast-TCO on ice for 30 min at 450 rpm. Later, the cells were centrifuged at 1500 rpm for 5 min at 4°C to remove unbound mAbs. Next, the cells were resuspended in PBS containing 1% w/v BSA and 0.1% w/v sodium azide. The radiolabeled antibodies (2.5 μg) [^89^Zr]Zr-DFO-Trast, [^89^Zr]Zr-DFO-Trast-Tz, or [^89^Zr]Zr-DFO-IgG-Tz were added to the cancer cells and incubated on ice during 90 min at 37 °C. PBS containing unbound radioimmunoconjugates was removed by centrifuging cells at 1500 rpm for 5 min at 4°C. The previous centrifugation steps were repeated three times and the activity bound to the cell pellet was measured on a gamma counter calibrated for ^89^Zr. The percent bound activity was calculated by dividing the cell pellet activity to the total activity in the suspension and cell pellet.

### Small animal PET imaging and biodistribution studies

Mice were first injected with TCO-conjugated mAbs (50 µg). At 24 h after injection of the TCO-conjugated mAbs, mice were injected with unconjugated or Tz-conjugated [^89^Zr]Zr-DFO-mAbs or [^64^Cu]Cu-NOTA-mAbs and used for both biodistribution and PET imaging studies. Acute biodistribution and PET imaging studies were performed at 48 h or 24 h after tail vein injection of the unconjugated or Tz-conjugated [^89^Zr]Zr-DFO-mAbs or [^64^Cu]Cu-NOTA-mAbs, respectively.

Mice used for biodistribution studies were sacrificed and organs were harvested, weighed, and assayed in a gamma counter. The radioactivity was expressed in each organ as a percentage of the injected dose per gram of the organ (%ID/g).

PET imaging was conducted on a on a Mediso nanoScan PET/CT scanner (Mediso) at 48 h or 24 h post-injection of [^89^Zr]Zr-DFO-mAbs or [^64^Cu]Cu-NOTA-mAbs modified with or without Tz. The mice were anesthetized by inhalation of 2% isofluorane (Baxter Healthcare) in an oxygen gas mixture before recording PET/CT images. PET/CT data for each group (n = 3) was recorded and images analyzed using 3D Slicer software (version 5.0.3, a free and open source software https://www.slicer.org/).

#### CD8^+^ T-cell imaging

CT26-hHER2 xenografts with a volume of ∼200 mm^3^, were randomly divided into control, no click and click groups. The mice in the control cohort received saline injection. In the no click and click groups, mice received the Per or Per-TCO (5 mg kg^-1^) injections on day 1. On day 2, mice in the no click or click groups received T-DXd or T-DXd-Tz (5 mg kg^-1^) injections. On day 3, [^64^Cu]Cu-NOTA-CD8 mAbs (mouse reactive, 2.5 mg kg^-1^, ∼7.4 Mbq) was intravenously injected in mice in the control, no click or click groups. The mice were then imaged at 24 and 48 h timepoints using the Mediso PET Scanner. The mice were anesthetized by inhalation of 2% isofluorane (Baxter Healthcare) in an oxygen gas mixture before recording PET/CT images. PET/CT data for each group (*n* = 3) was recorded. Image analyses and regions of interest (ROI) quantifications were done using 3D Slicer software (version 5.0.3, a free and open source software https://www.slicer.org/).

### *In vivo* therapeutic efficacy studies

All animal experiments were conducted in accordance with guidelines approved by Washington University School of Medicine’s Research Animal Resource Center and Institutional Animal Care and Use Committee. Eight- to 10-week-old *nu/nu* female or male mice and 4 to 6-week-old Balb/c female or male mice were obtained from Charles River Laboratories. The therapies were initiated when the tumor volume reached ∼200 mm^3^. Tumor volumes were measured twice a week by caliper measurements. Humane endpoints were defined as tumor volumes >1,500 mm^3^, bodyweight loss of >20%, tumor ulceration, and other clinical symptoms of acute toxicity. At the terminal stage, tumors were collected for the western blot analyses.

#### HER2^+^ gastric cancer model

Mice bearing HER2^+^ NCIN87 xenografts were stratified into the following groups: control (saline), Panitumumab (Pan), T-DXd, no click (Panitumumab plus T-DXd) and click (Pan-TCO plus T-DXd-Tz) cohort. Pan-TCO or Pan was administered at 10 mg kg^-1^ *via* tail vein. At 24 h post-injection of Pan-TCO, the mice received an intravenous injection of T-DXd or T-DXd-Tz (10 mg kg^-1^). Mice in Pan or T-DXd cohorts received intravenous injection of Pan or T-DXd (10 mg kg^-1^) alone.

#### HER2^+^/HER2^-^ bilateral tumor model

*Nu/nu* female or male mice (Charles River Laboratories) were injected subcutaneously with 5 million NCIN87 cells, and/or 0.25 million A431 cells in a 100 μL cell suspension of a 1:1 (v/v) mixture of medium with reconstituted basement membrane (BD Matrigel, BD Biosciences). Mice-bearing NCIN87/A431 bilateral tumors were developed by subcutaneous injection of NCIN87 and A431 cancer cells on the bilateral dorsal flank regions. When the tumor volume reached the size of 200 mm^3^, mice were randomized them into six cohorts: control (saline), Pan (5 mg kg^-1^), ADC T-DXd alone (5 mg kg^-1^), no click (Pan plus T-DXd, both at 5 mg kg^-1^), and click (Pan-TCO plus T-DXd-Tz, both at 5 mg kg^-1^).

#### NCIN87 gastric cancer model of acquired resistance to ADC

*Nu/nu* male and female mice bearing NCIN87 xenografts were treated with T-DXd monotherapy intravenously (5 mg kg^-1^). Nearly one month after therapy, mice were stratified into responders *versus* non-responders, based on the tumor volume measurements, western blot analyses, and HER2-PET imaging. HER2-targeting immuno-PET was performed at 24 h post-intravenous injection of [^64^Cu]Cu-NOTA-trastuzumab (50 µg, 200 µCi). The mice that initially did not respond to T-DXd therapy (*i.e.,* non-responder) were treated intravenously with Pan-TCO (5 mg kg^-1^) on day 1 and T-DXd-Tz on day 2 (5 mg kg^-1^ as described above), while the responders continued to receive T-DXd therapy (5 mg kg^-1^).

#### BT474-trastuzumab resistant model

*Nu/nu* female mice were implanted subcutaneously with one million estrogen receptor-positive BT474-trastuzumab resistant cells. Drinking water of mice was supplemented with 0.67 μg mL^−1^ of β-estradiol (Sigma) from 1 week in advance of tumor inoculation and continued until mice were sacrificed. Fresh-estradiol supplemented water was provided twice a week. The T-DXd ADC therapy was initiated when the tumor volume reached 200 mm^3^. The tumor volumes were measured twice a week for a period of ∼1 month. Mice were then stratified into responders versus non-responders, based on the tumor volume measurements. The responder mice were maintained on T-DXd (5 mg kg^-1^) therapy and non-responder mice on the click therapy. In the click therapy cohort, the mice were injected with Per-TCO mAbs (5 mg kg^-1^) on day 1. Following this, the mice were injected with T-DXd-Tz (5 mg kg^-1^) on day 2 and tumor volumes were measured twice a week.

#### Immunocompetent CT26-hHER2 model

CT26-hHER2 cancer cells (0.25 million) were xenografted on the right shoulder of female BALB/c mice and treatments were initiated when the tumor volumes reached the size of 200-500 mm^3^. Then, mice were randomized into six cohorts: control (saline), Per (5 mg kg^-1^), T-DXd (5 mg kg^-1^), no click (Per plus T-DXd, 5 mg kg^-1^), and click (Per-TCO plus T-DXd-Tz, 5 mg kg^-1^). The mice that showed complete remission of the tumors after the therapy, were further rechallenged with 5 million of CT26-hHER2 cancer cells.

### Statistical analyses

Data were analyzed using GraphPad Prism software version 10.00 (www.graphpad.com). Statistical comparisons of mean values between experimental groups were performed using analysis of variance (ANOVA) followed by Student’s t-tests. A *P*-value less than 0.05 was considered statistically significant.

To investigate whether changes in tumor volume were significant in the different cohorts, least square means for each effect were estimated. All analyses were conducted using SAS 9.4 statistical software at the two-sided 5% significance level.

## Acknowledgments

We thank the Washington University School of Medicine isotope production team for the production of zirconium-89 and copper-64, the small animal imaging facility for help with the small animal PET/CT image collection, and the Washington University Center for Cellular Imaging (WUCCI) for their assistance with the TEM samples preparation and image acquisition. We thank Dr. Katherine Basore for her help with the TEM image analyses. We thank Dr. Luis Batista for letting us use their Odyssey CLX Imaging System. We thank Dr. Luke Carter for his help with the 3D Slicer software. We thank Dr. Ricardo D’Oliveira Albanus for his help with data presentation and figure preparation.

## FIGURE LEGENDS – EXTENDED FIGURES

**Extended Figure 1:**
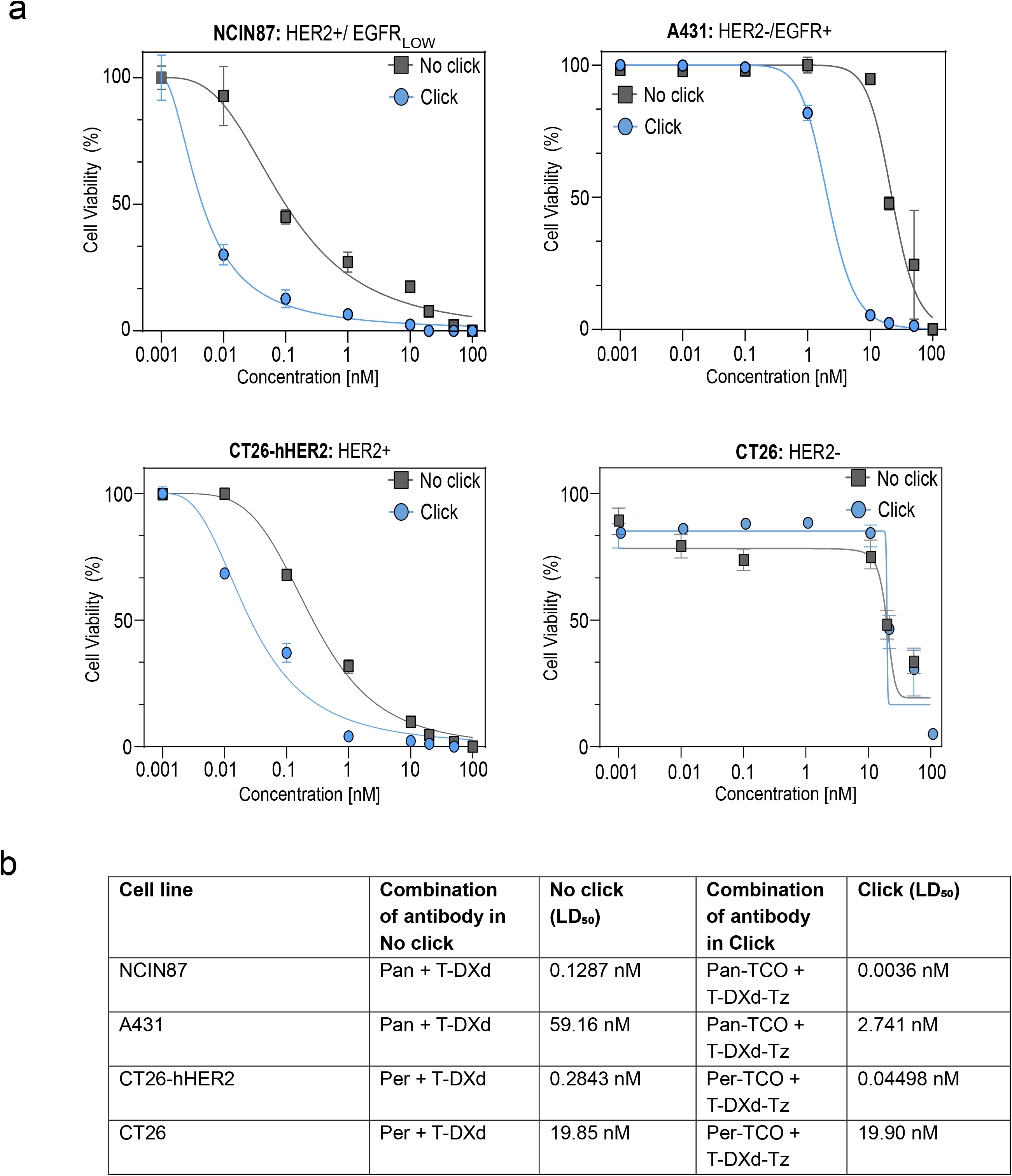
Mass spectrometry to quantify TCO or Tz click moieties per antibody. **(a)** Schematic representation of the mAbs conjugated to trans-cyclooctene-(PEG)_4_-NHS (TCO) or Tetrazine-(PEG)_5_-NHS (Tz) with a molar ratio of 15click moieties per mAbs. **(b)** MALDI spectrum of the mAbs, antibody-TCO and antibody-Tz. **(c)** Table shows quantification of TCO or Tz click moieties per mAbs.

**Extended Figure 2:**
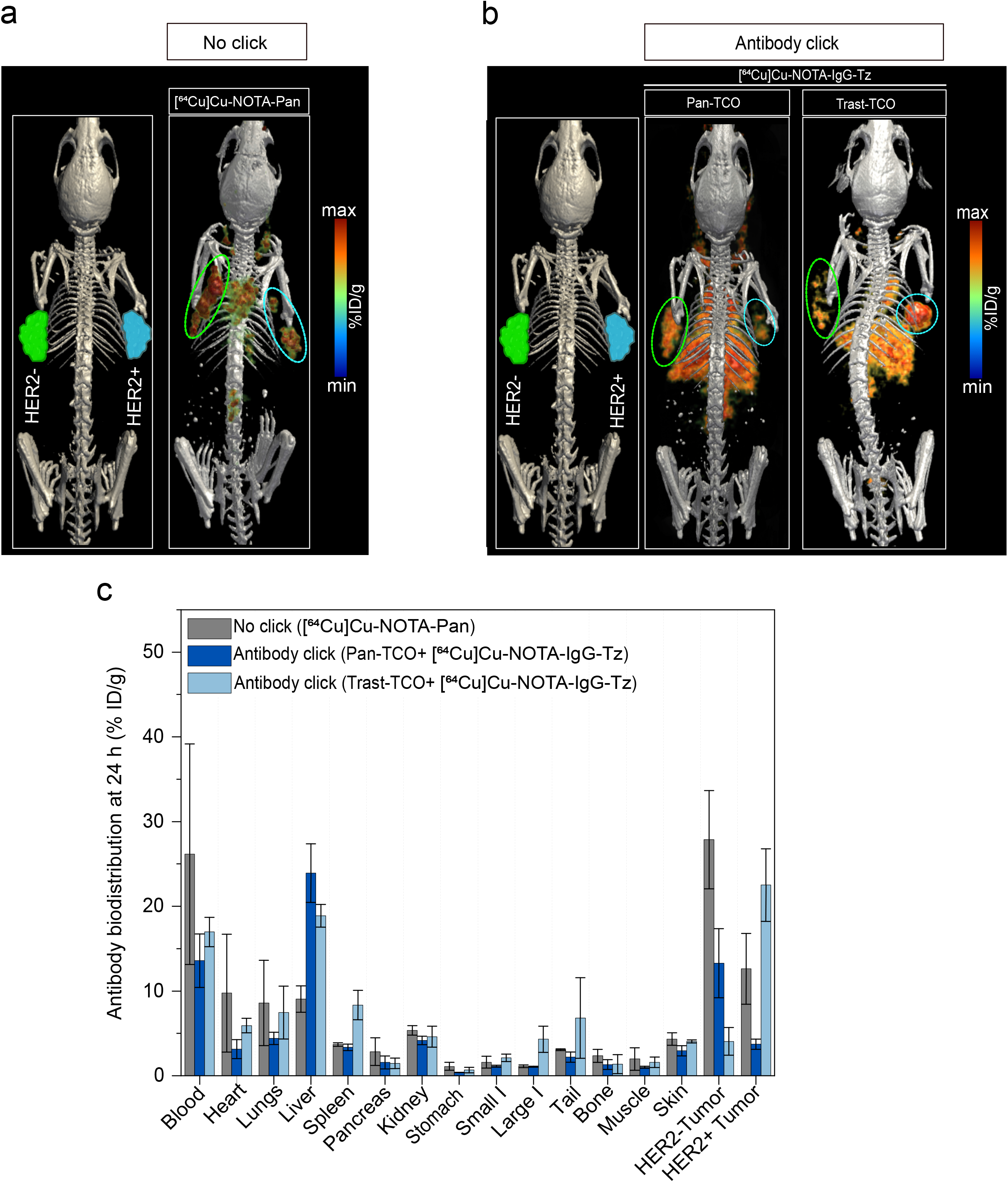
Antibody click improves the uptake and internalization rate. **(a)** Immunofluorescence analyses of the click mAbs *versus* no click mAbs. The fluorescence intensity was obtained by quantitative analyses of fluorescence in A431, BT474 trastuzumab resistant, CT26-hHER2, CT26 cancer cells cultured on 96 well plates. The combination of antibodies used for the different cell lines are depicted in table. **(b)** Fluorescence microscopy images of the no click and click groups with mAbs labeled with the pH-sensitive dye pHrodo, which fluoresces upon internalization. The combination of antibodies used for different cell line are depicted in the table.

**Extended Figure 3:**
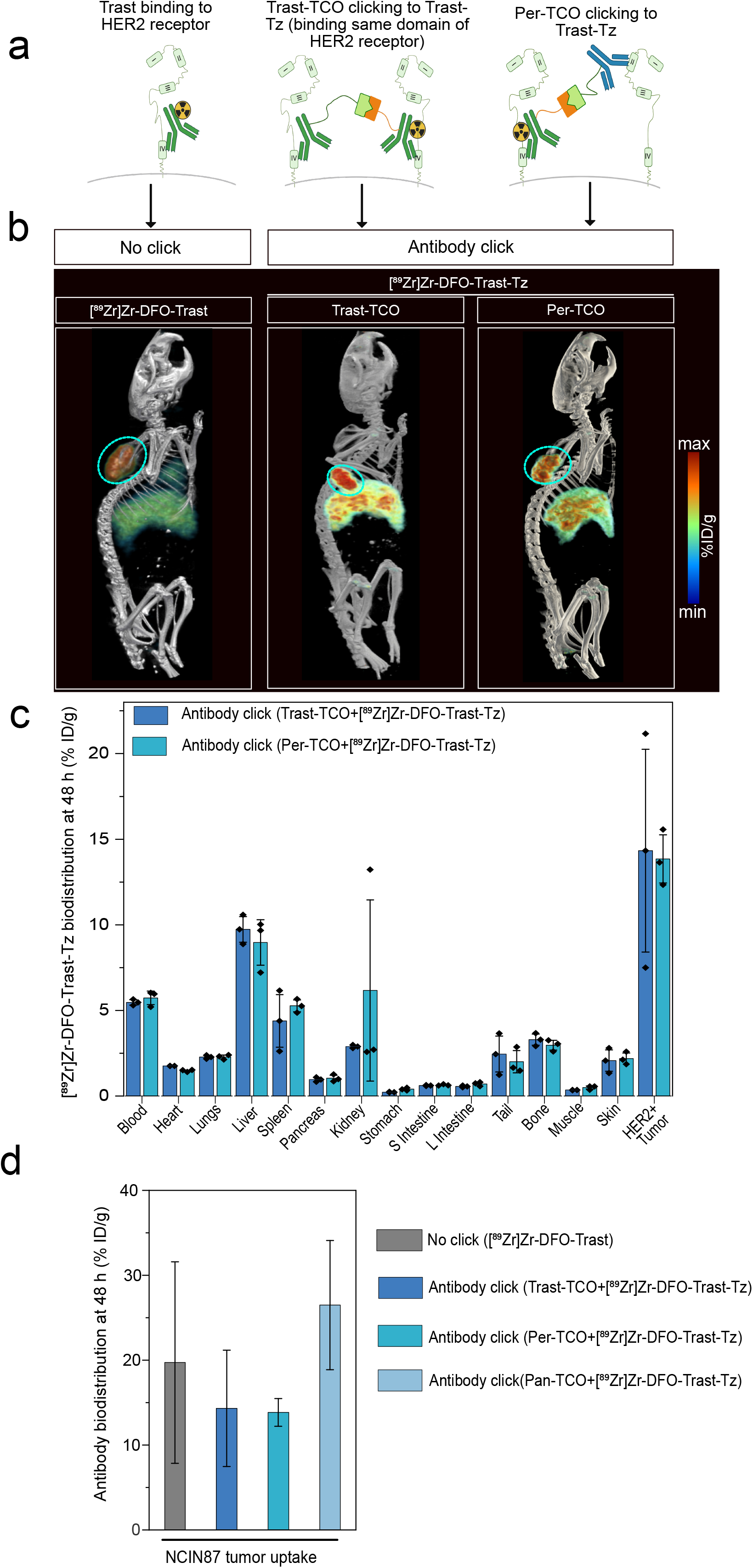
Immuno-PET of no click versus click mAbs. **(a)** Schematic representation of the trastuzumab and pertuzumab binding site on the domain IV and domain II of HER2 extracellular protein, respectively. **(b)** Representative PET maximum intensity projection (MIPs) image of the mice bearing the NCIN87 (HER2^+^/EGFR-low) tumor injected with the no click and clicking antibodies and imaged at 48 h timepoint (n=3). **(c)** Biodistribution data at 48 h post-injection of clicking mAbs. The first pair of the clicking antibodies *i.e*., Pan-TCO (50 µg) was injected via the tail vein 24 h prior to the [^89^Zr]Zr-DFO-Trast-Tz (50 µg, ∼7.4 Mbq). Biodistribution was performed at 48 h post injection. Bars, n=3 mice per group, mean ± S.E.M. %ID g^−1^, percentage of injected dose per gram. **(d)** Comparative tumor uptake analyses of the no click and different click antibodies combination in the mice bearing the NCIN87 tumors at 48 h timepoint.

**Extended Figure 4:**
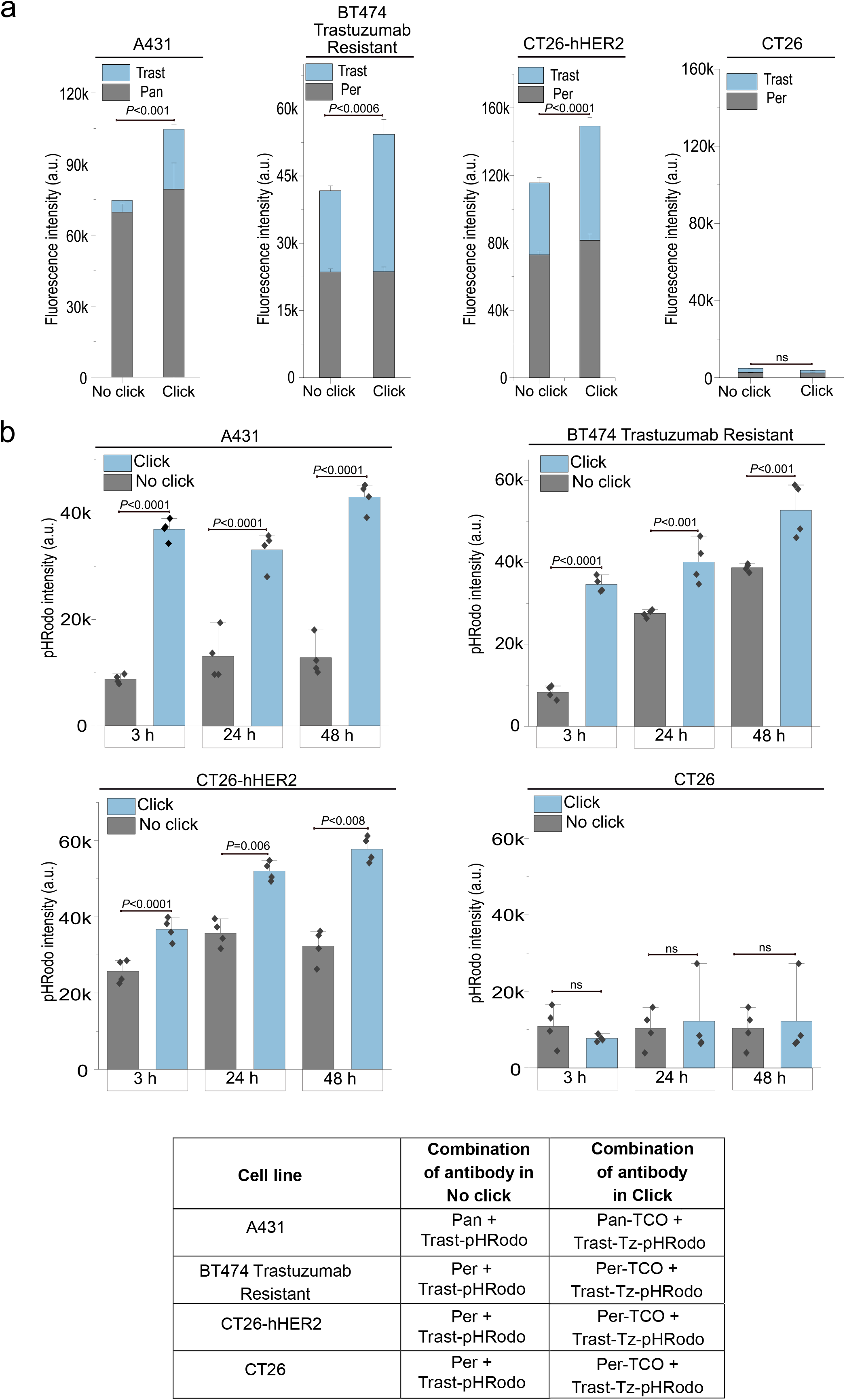
Pan-TCO and Trast-TCO clicking with IgG-Tz confirms the clicking at tumor surface. (a-c) Representative PET MIPs image of a bilateral tumor model: NCIN87 tumors (HER2^+^/EGFR-low) on the right flank and A431 tumors (HER2^-^/EGFR^+^) tumors on the left flank. The first pair of click antibodies *i.e.,* Pan-TCO or Trast-TCO was injected *via* the tail vein 24 h prior to the injection of [^64^Cu]Cu-NOTA-IgG-Tz (50 µg, ∼7.34 Mbq). The no click mice were intravenously injected with [^64^Cu]Cu-NOTA-Pan (50 µg, ∼7.4 Mbq) and imaged at 24 h after injection. All the mice were imaged at 24 h timepoint followed by a biodistribution study. Bars, n=3 mice per group, mean ± S.E.M. %ID g^−1^, percentage of injected dose per gram.

**Extended Figure 5:**
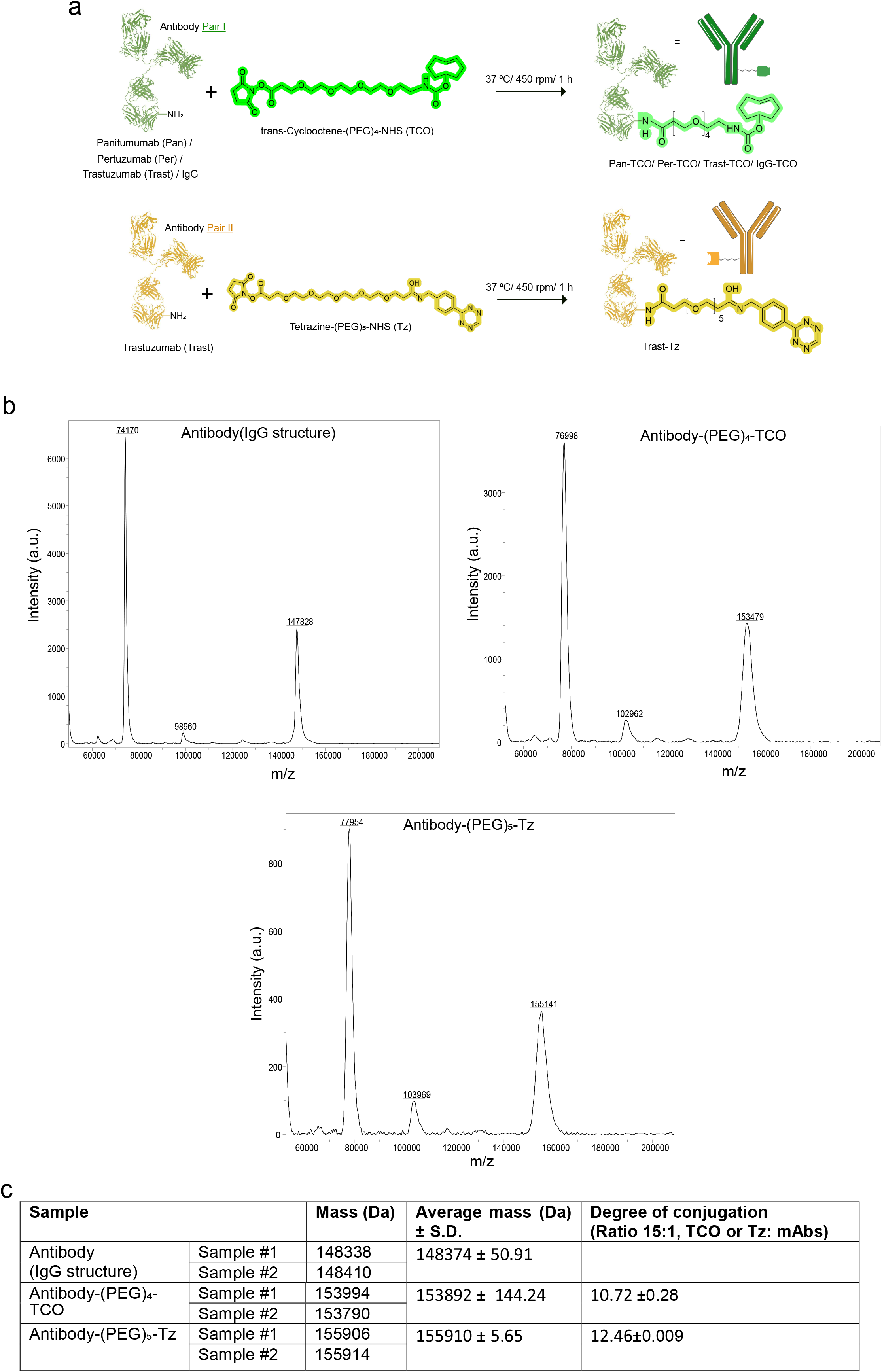
Antibody click improves the uptake of ADC irrespective of the presence of its binding receptor. Cell viability (WST-8) assay comparing no click versus click conditions to validate the efficiency of ADC delivery in NCIN87, A431, CT26-hHER2 and CT26 (control) cell lines. The table shows the different antibody-ADC combinations tested on each cell line.

