## Supplementary material for "In Vivo Biorthogonal Antibody Click for Dual Targeting and Augmented Efficacy in Cancer Treatment": SI file

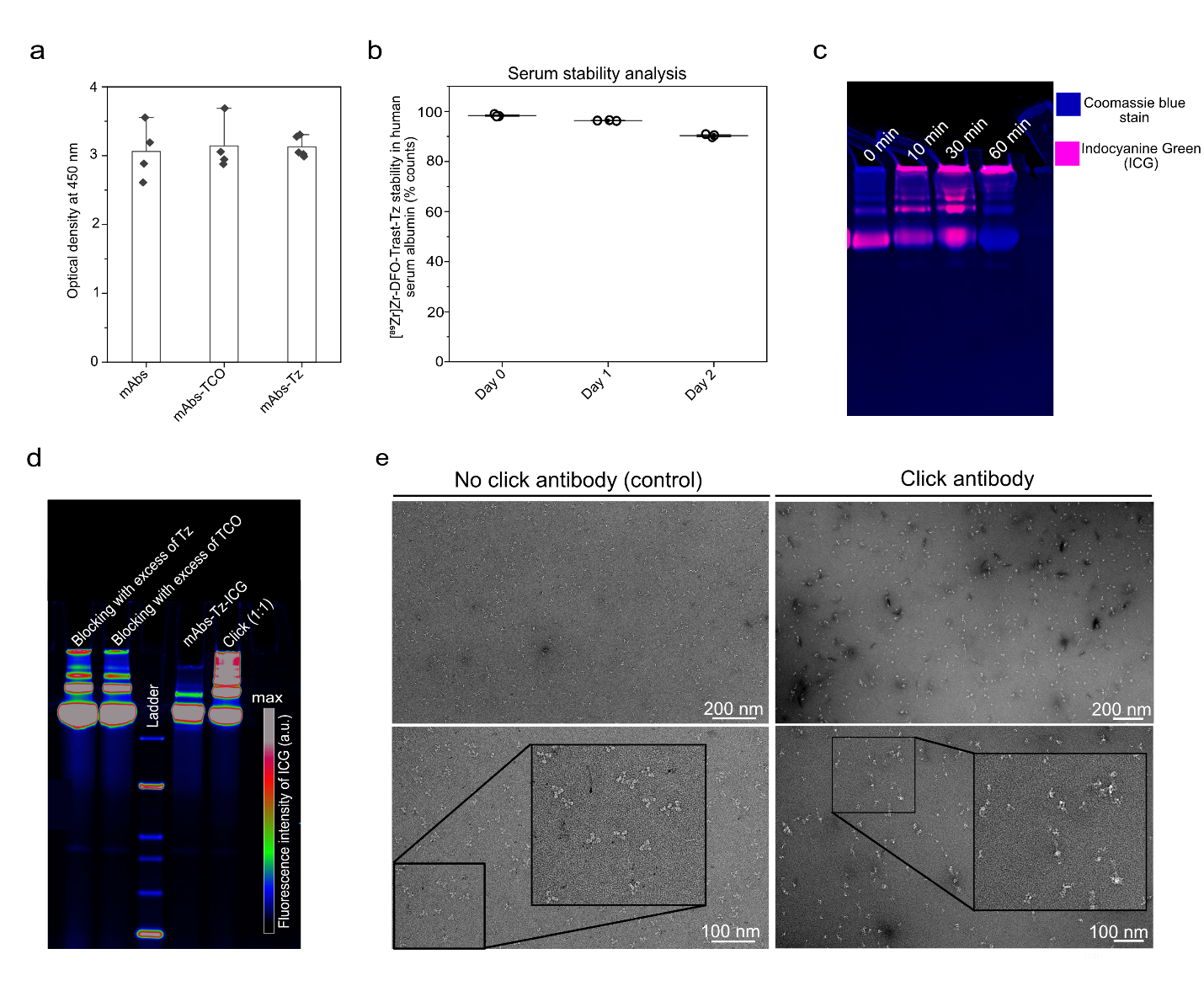

**Supplementary Figure 1.**

**a)** Immunoreactivity of the immunoconjugates (i.e, Trast, Trast-TCO, and Trast-Tz) to the HER2 receptor protein.

**b)** Assessment of the stability of the [^89^Zr]Zr-DFO-Trast-Tz in the human serum albumin (HSA) by instant thin layer chromatography (iTLC) (n=3).

**c)** SDS-PAGE analyses and quantification of the clicking antibodies at a 1:1 ratio over different incubation times.

**d)** SDS-PAGE analyses of the click reaction blocked with an excess of the unconjugated TCO or Tz.

**e)** Negative-staining-Transmission Electron Microscopy (TEM) images of the no-click mAbs versus clicking mAbs. Clicking mAbs were prepared by reacting mAbs-TCO and mAbs-Tz in a 1:1 ratio for 90 min.

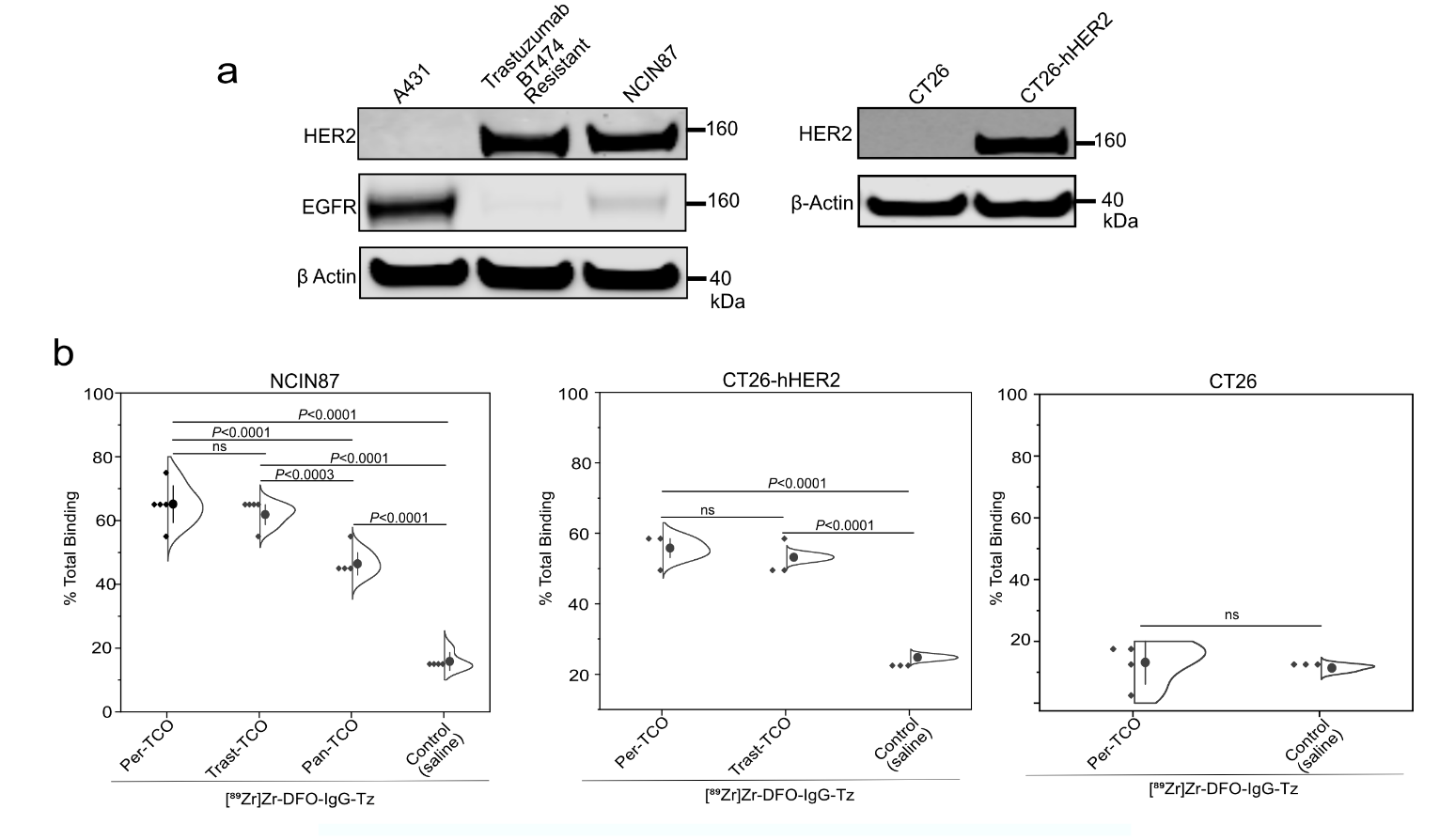

**Supplementary Figure 2.**

**a)** Western blot analysis of the different cells expressing tHER2 and EGFR.

**b)** Radio-immunoreactivity assay of the mAbs-TCO bound to the cellular surface reacting with the [^89^Zr]Zr-DFO-IgG-Tz.

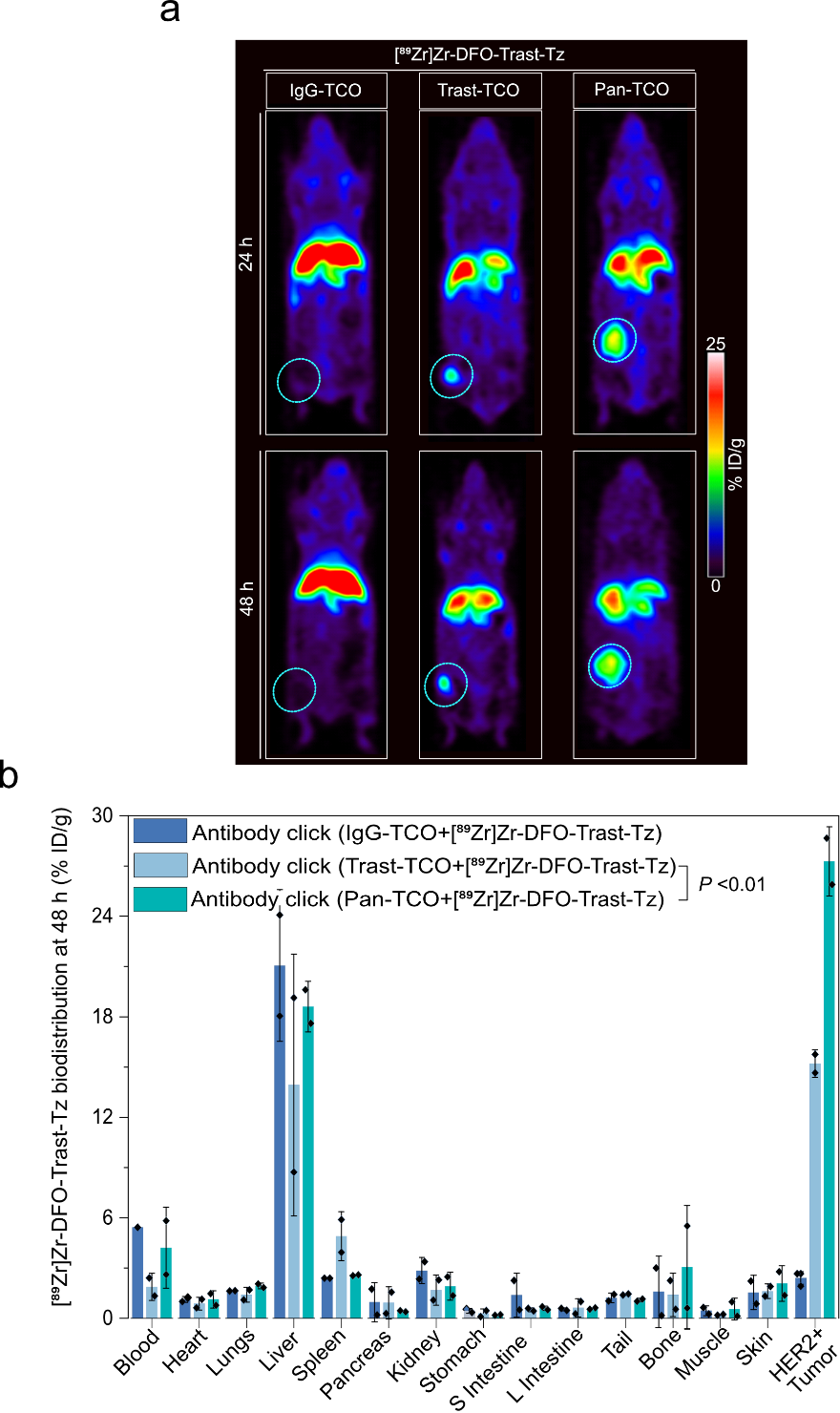

**Supplementary Figure 3*.***

**a)** Representative coronal PET images of the mice bearing the NCIN87 (HER2+/EGFR-low) tumor injected with the clicking antibodies and imaged at 48 h time point.

**b)** Biodistribution was performed at 48 h post-injection. Bars, n=2 mice per group, mean ± S.E.M. %ID g^−1^, percentage of injected dose per gram.

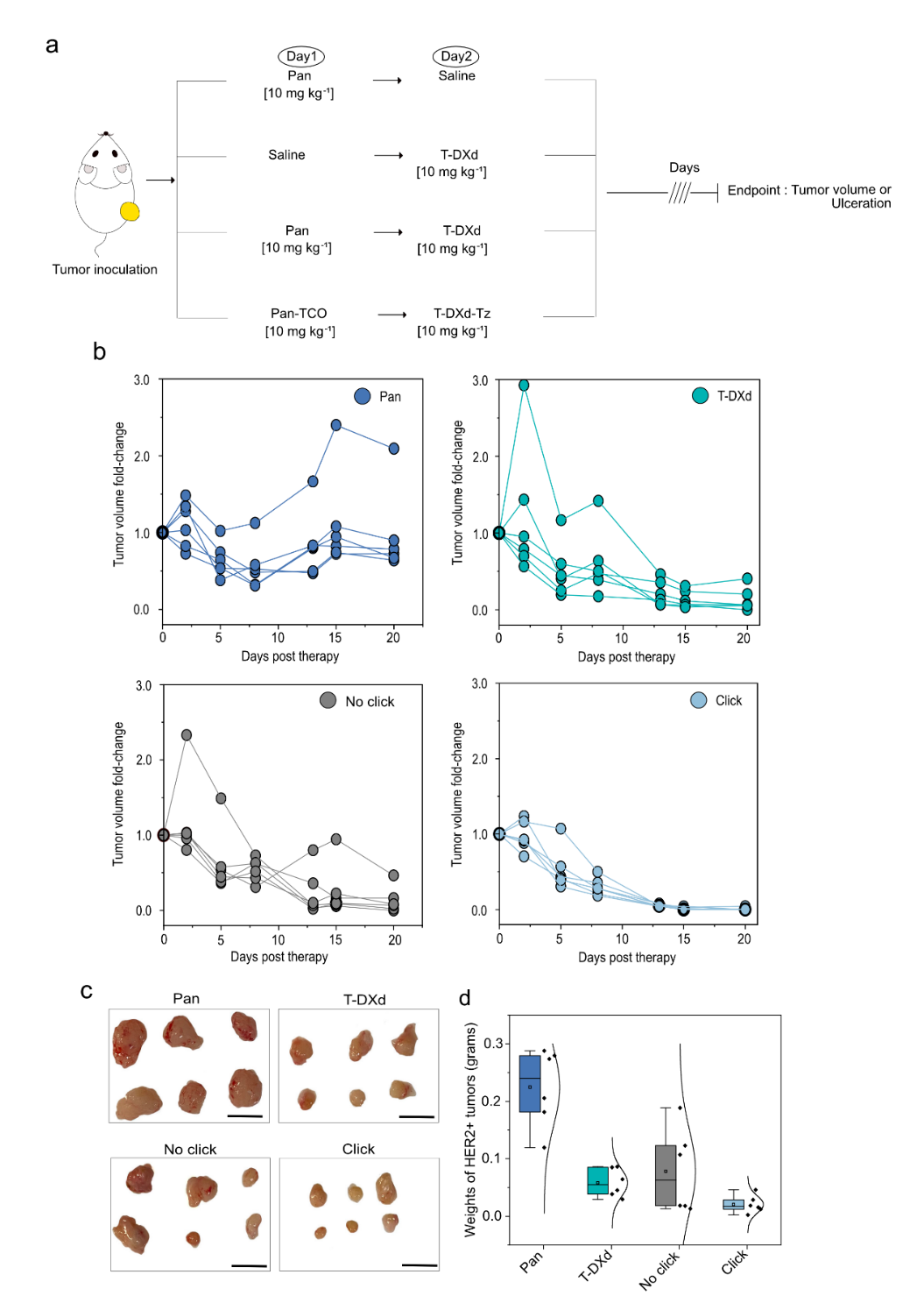

**Supplementary Figure 4.**

**a)** Schematic representation of the NCIN87 (HER2+) cancer tumor model stratified into different cohorts: Pan, T-DXd, no click (Pan plus T-DXd), and antibody-ADC click (Pan-TCO + T-DXd-Tz).

**b)** Represents the in vivo therapeutic efficacy of different cohorts after receiving a single dose of intravenous injection of each antibody started at day 0. For the mice receiving the combination of antibodies, the first pair of click antibodies i.e., Pan or Pan-TCO (10 mg kg^-1^) was injected via the tail vein 24 h prior to the injection of T-DXd or T-DXd-Tz (10 mg kg^-1^). Bars, n=6 mice per group, mean ± standard error.

**c-d)** Photographic images and quantitative analysis of the tumor excised on twenty-sixth-day post-therapy, respectively.

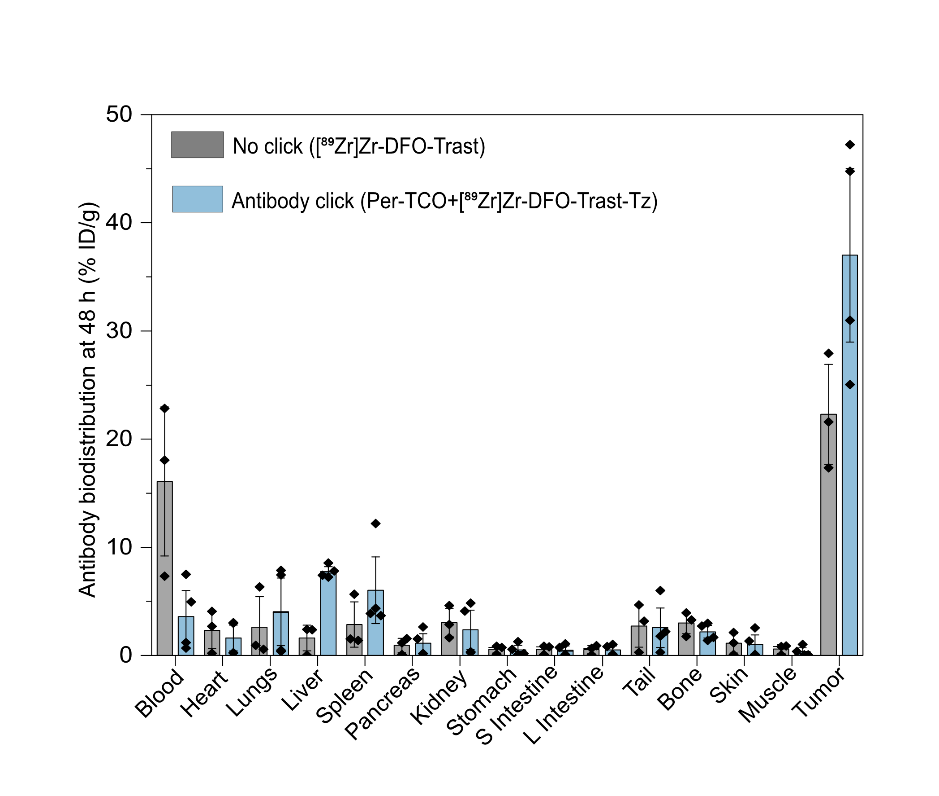

**Supplementary Figure 5.** Biodistribution analysis of the click versus no click antibodies in the immunocompetent mice performed at 48 h post-injection. The first pair of click antibodies i.e., Per-TCO was injected via the tail vein 24 h prior to the injection of [^89^Zr]Zr-DFO-Trast-Tz (50 µg, ~7.4 Mbq). Bars, n=3 mice per group, mean ± S.E.M. %ID g^−1^, percentage of injected dose per gram.

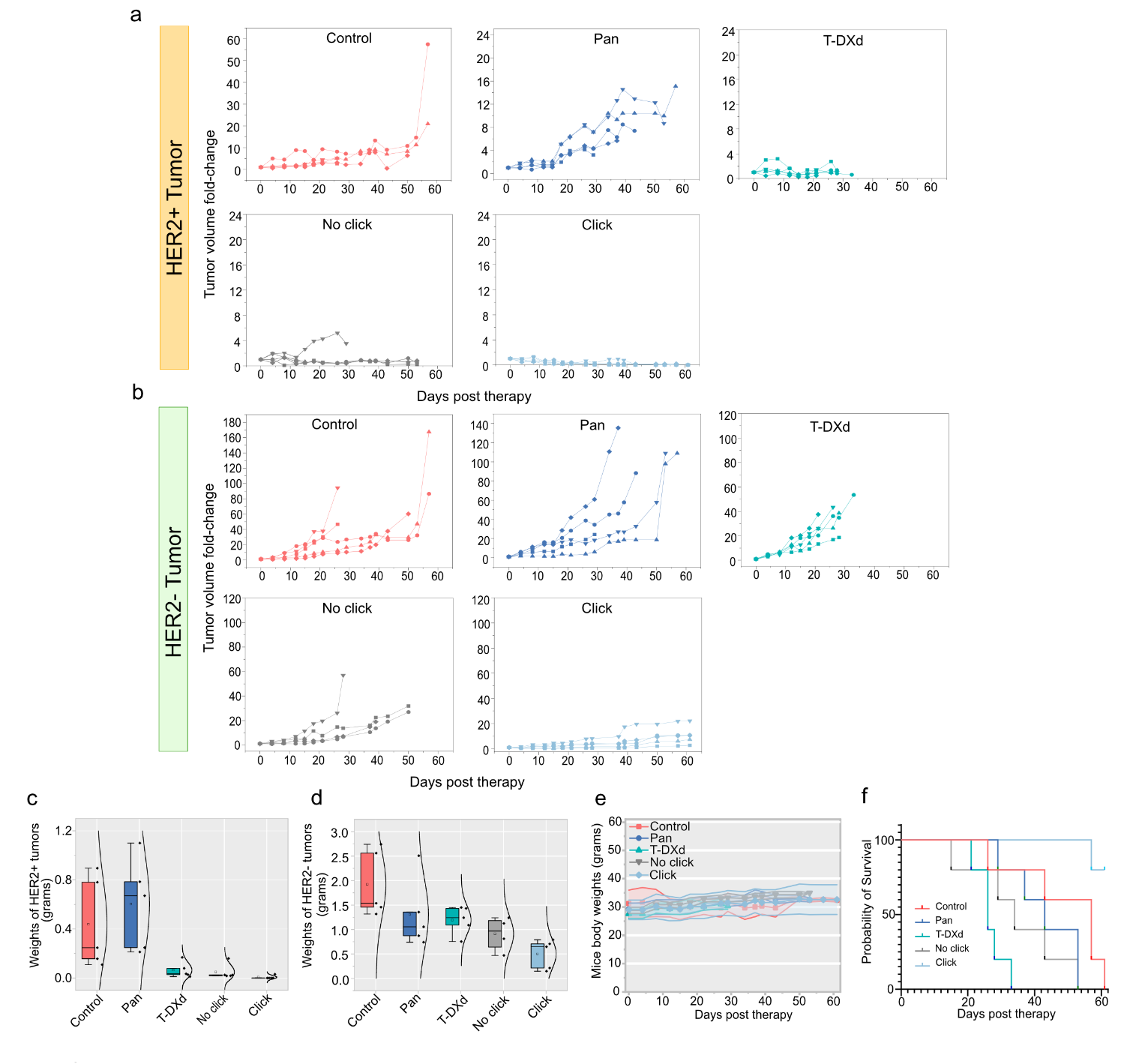

**Supplementary Figure 6.**

**a)** In vivo therapeutic efficacy of different cohorts after receiving a single dose of intravenous injection of each antibody was started at day 0 in the bilateral tumor model. For the mice receiving the combination of drugs, the first pair of click antibodies i.e., Pan or Pan-TCO (10 mg kg^-1^) was injected via the tail vein 24 h prior to the injection of T-DXd or T-DXd-Tz (10 mg kg^-1^). Bars, n=5 mice per group, mean ± standard error.

**c-d)** Quantitative analysis of HER2- and HER2+ tumor excised post-therapy, respectively. **e)** Body weight of the bilateral tumor bearing mice post-therapy. **f)** Median survival of bilateral tumor bearing mice post-therapy.

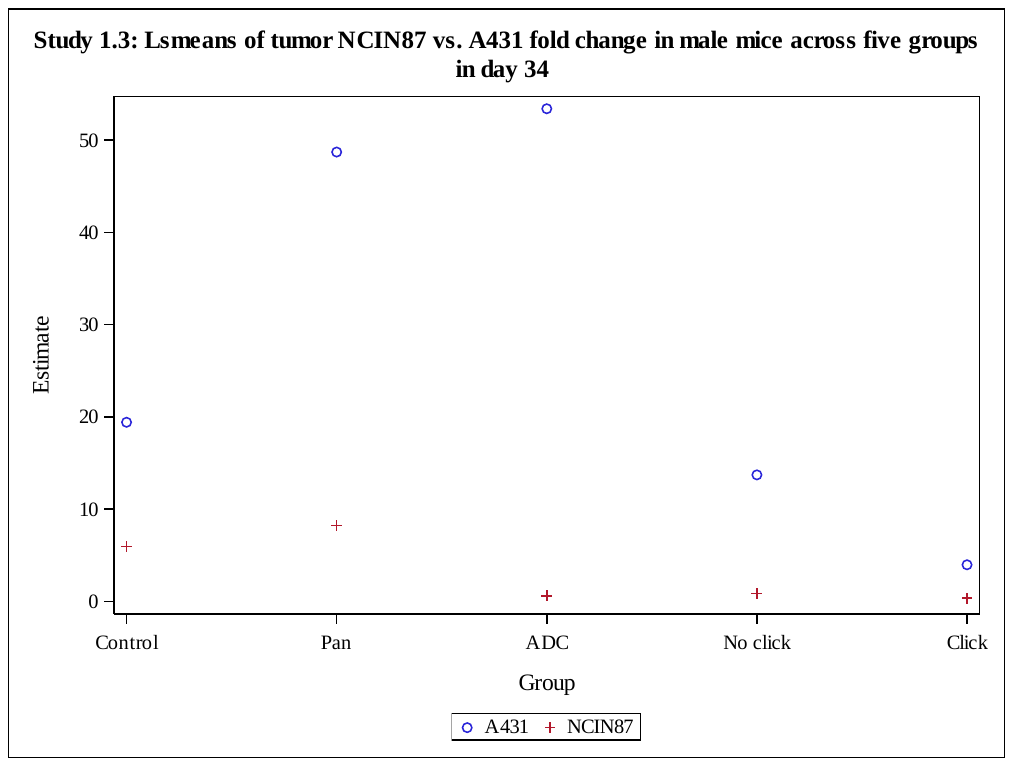

**Supplementary Figure 7.**

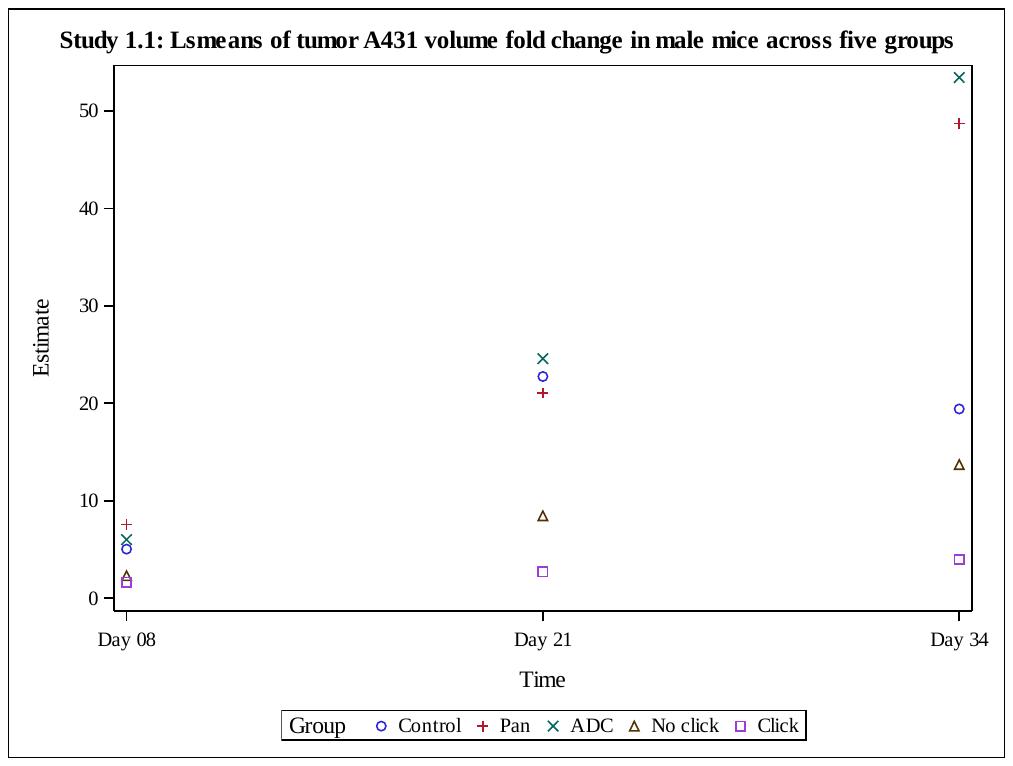

**Supplementary Figure 8.**

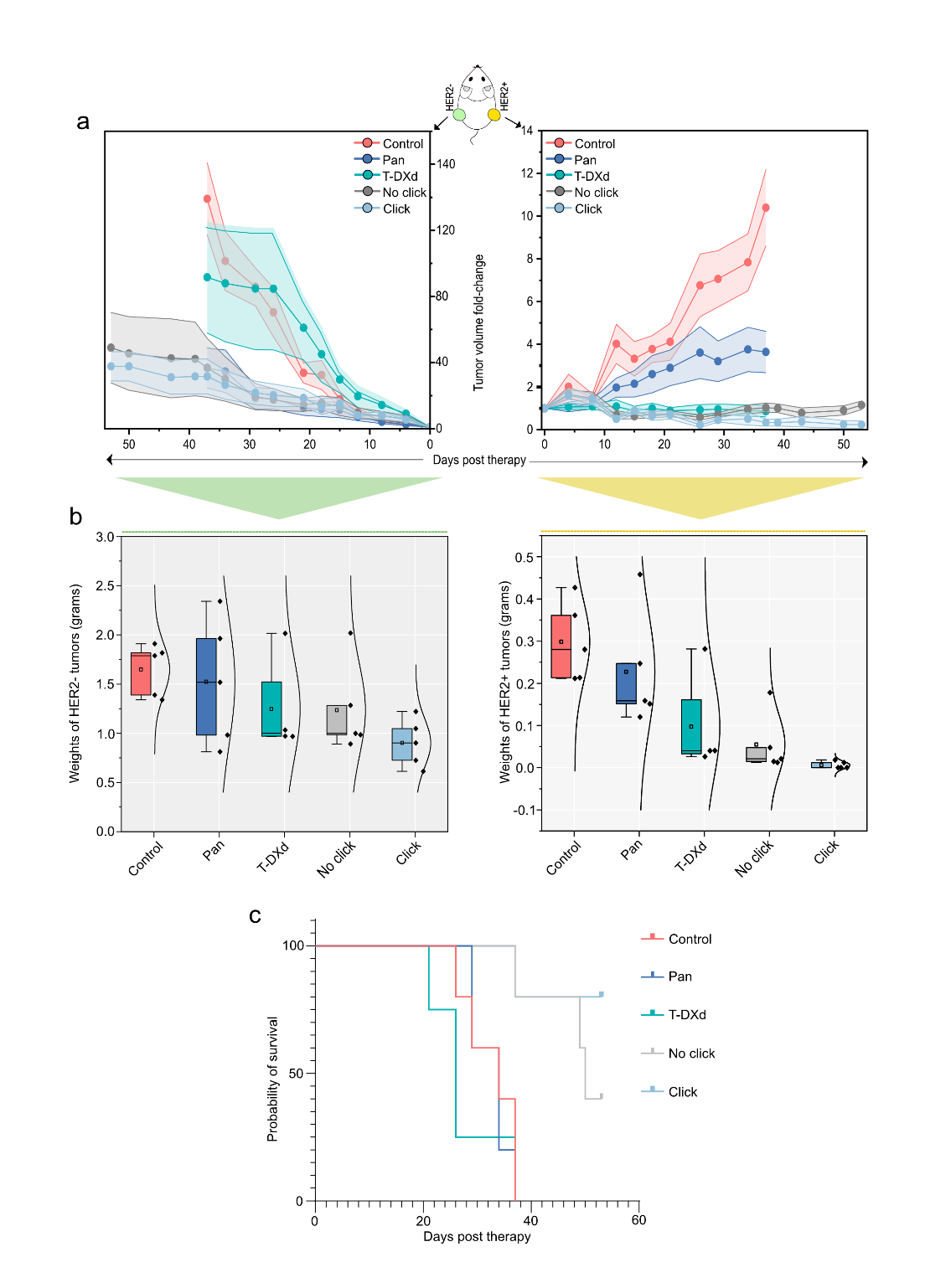

**Supplementary Figure 9.**

**a)** In vivo therapeutic efficacy of different cohorts (i.e., control saline, Pan, no click Pan+ T-Dxd and antibody-ADC click Pan-TCO plus T-DXd-Tz )validated in female bilateral tumor-bearing mice. For the mice receiving the combination of drugs, the first pair of click antibodies i.e., Pan or Pan-TCO (5 mg kg^-1^) was injected via the tail vein 24 h prior to the injection of T-DXd or T-DXd-Tz (5 mg kg^-1^). Bars, n=5 mice per group, mean ± standard error.

**b)** Quantitative analysis of HER2- and HER2+ tumor excised post-therapy, respectively.

**c)** Median survival of bilateral tumor-bearing mice post-therapy.

Day 07/08

Day 15

Day 21/22

Day 29/30

Time

0

20

40

60

Estimate

Click

No click

ADC

Pan

Control

Group

Study 1.1: Lsmeans of tumor A431 volume fold change across five groups

**Supplementary Figure 10.**

Day 07/08

Day 15

Day 21/22

Day 29/30

Time

0

2

4

6

Estimate

Click

No click

ADC

Pan

Control

Group

Study 1.2: Lsmeans of tumor NCIN87 volume fold change across five groups

**Supplementary Figure 11.**

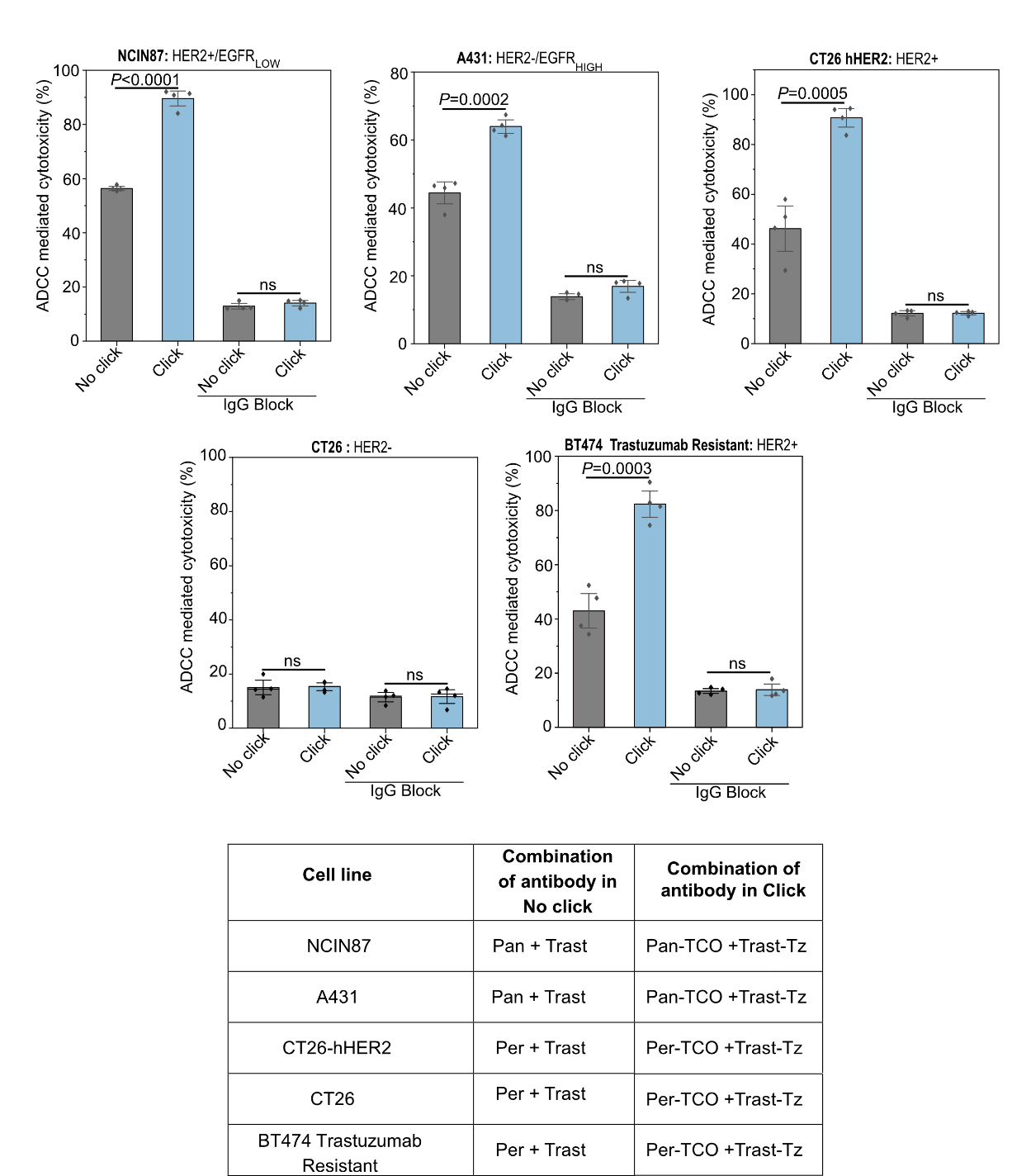

**Supplementary Figure 12.** ADCC LDH assay (cytotoxicity) response to click and no click antibody treated to target NCIN87, A431, CT26 hHER2, CT26, and BT474 trastuzumab-resistant cancer cells. Target cells were treated with a combination of the antibodies depicted in the table at a 1:1 ratio followed by the addition of human PBMCs. After 48 h of the addition of human PBMCs, the LDH assay was performed (LDH-based readout). Bars, n=4, mean ± standard deviation; data were analyzed using unpaired T-test using GraphPad Prism.

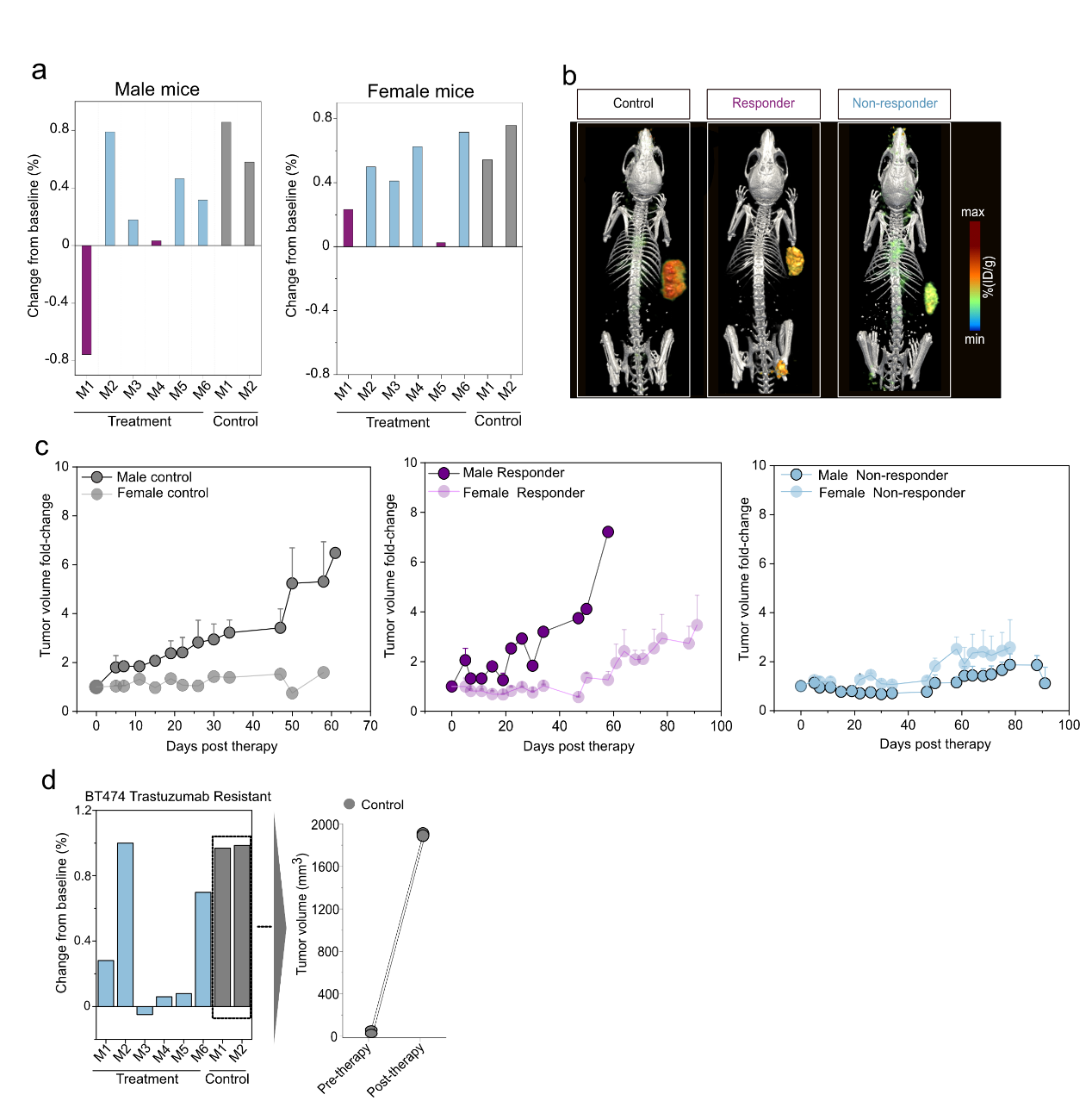

**Supplementary Figure 13.**

**a)** Represents the baseline change in the NCIN87 tumor volume of male and female mice after receiving monotherapy of T-DXd.

**b)** Anti-HER2 PET imaging to monitor HER2 protein levels in the responder versus non-responder tumors in female mice upon stratification on day 30. The mice were intravenously injected with [^64^Cu]Cu-NOTA-Trast-Tz (80 µg, ~10.5-11.84 Mbq), and PET imaging was performed at 24 h timepoint (n=3).

**c)** Tumor volume measurements of the male and female mice in the saline-treated versus mice responding and non-responding to T-DXd monotherapy (5 mg kg^-1^).

**d)** Represents the change in the BT474 trastuzumab-resistant tumor volume after receiving monotherapy of T-DXd.

Non-responder

Responder

Group

0.8

0.9

1.0

1.1

1.2

Estimate

Male

Female

Gender

Study 2.2: Lsmeans of volume fold change between responder and non-responder

(

gender effect

)

**Supplementary Figure 14.**

**Supplementary Table 1.** Cancer cell lines and media composition used in this work.

| **Cell line/Tumor type** | **Media Composition** |
| --- | --- |
| NCIN87/gastric cancer | RPMI-1640 growth medium + 10% fetal calf serum (FCS) + 2 mM L-glutamine + 10 mM hydroxyethyl piperazineethanesulfonic acid (HEPES), 1 mM sodium pyruvate + 4.5 g L^−1^ glucose + 1.5 g L^−1^ sodium bicarbonate + 1% of penicillin (10000U) and 10 mg mL^−1^ of streptavidin. |
| CT26/colorectal cancer | (RPMI)-1640 growth medium + 10% fetal calf serum (FCS) + 2 mM L-glutamine + 1% of penicillin (10000U) and 10 mg mL^−1^ of streptavidin. |
| A431/epidermoid cancer | DMEM with high glucose + 10% fetal calf serum (FCS) + 4mM L-Glutamine + 1mM Sodium Pyruvate + 4.5 g L^−1^  Glucose + 1.5 g L^−1^  Sodium Bicarbonate + 1% of penicillin (10000U) and 10 mg mL^−1^ of streptavidin. |
| BT474/breast cancer | Dulbecco’s modified Eagle medium: Nutrient Mixture F-12 (DMEM/F-12) + 10% fetal calf serum (FCS) + 5 mL of non-essential amino acid (100X) + 2mM glutamine + glucose + 1% of penicillin (10000U) and 10 mg mL^−1^ of streptavidin. |

**Supplementary Table 2.** FDA-approved antibodies and ADCs used in this work.

| **Antibody** | **Antibody-drug conjugate** | **Target** |
| --- | --- | --- |
| Panitumumab |  | EGFR |
| Pertuzumab |  | HER2, domain II |
| Trastuzumab |  | HER2, domain IV |
|  | T-DM1 | HER2, domain IV |
|  | T-DXd | HER2, domain IV |
